# Optimal transport modeling uncovers spatial domain dynamics in spatiotemporal transcriptomics studies

**DOI:** 10.1101/2025.10.06.680672

**Authors:** Wenjing Ma, Siyu Hou, Lulu Shang, Jiaying Lu, Xiang Zhou

## Abstract

Spatiotemporal transcriptomics is an emerging and powerful approach that adds a temporal dimension to spatial transcriptomics, enabling the characterization of dynamic changes in tissue architecture during development or disease progression. Here, we present SpaDOT (**Spa**tial **Do**main **T**ransition detection), a computational method designed to identify spatial domains and infer their temporal dynamics across time points for spatiotemporal transcriptomics studies. SpaDOT employs a variational autoencoder (VAE) framework to capture a low-dimensional representation of the data, incorporating hidden clustering variables to define distinct spatial domains. In the process, SpaDOT integrates a Gaussian Process prior and graph neighbor information to explicitly model both global spatial continuity and local structural heterogeneity among spatial locations within each time point, while using optimal transport (OT) to derive time-varying embeddings and infer the relationships between spatial domains across time points. Through simulation and real data applications, we demonstrate the superiority of SpaDOT in spatial domain detection, latent space preservation, and domain transition tracking over time. In real data applications, SpaDOT accurately captures dynamic domain transitions, including the disappearance and re-emergence of domains due to technical variation. Notably, SpaDOT uncovers key aspects of valvulogenesis in the developing heart, revealing the splitting of a single valve structure into two distinct functional valves -- the atrioventricular valve and the semilunar valve.

## Introduction

In recent years, spatial transcriptomics (ST) techniques have enabled the measurement of gene expression along with spatial coordinates within tissues, revolutionizing the field of genome biology^1^. These ST techniques can be broadly categorized into two main groups based on experimental protocols: *in situ* hybridization-based approaches, such as MERFISH^2^, seqFISH^3^, STARmap^4^, and 10X Xenium^5^; and next-generation sequencing-based approaches, such as 10X Visium^6^, Slide-seq^7^, Stereo-seq^8^. These techniques vary in spatial resolution and gene coverage, and collectively, they have been widely applied to uncover cellular heterogeneity^9,10^, map cell-cell interactions^11–13^, detect functional domains and microenvironments^10,14,15^, and characterize spatial gene expression patterns^16–19^. The application of ST techniques has provided critical insights into disease progression, including tumor evolution and neurodegenerative disorders, facilitating the development of spatially targeted therapies for disease treatments^20,21^.

An important analytic task in spatial transcriptomics is identifying spatial domains within complex tissues. Spatial domains are regions within a tissue characterized by relatively similar gene expression profiles, often corresponding to distinct biological functions. They serve as the fundamental architectural units of tissues, organizing different cell types to coordinate their roles in development, homeostasis, and disease progress. Accurately detecting these domains is therefore crucial for understanding tissue functions. Several computational methods have been developed for this purpose. For instance, methods like BASS^14^ and IRIS^1^ inferred spatial domains based on cell type compositions, while approaches such as SpatialPCA^15^, spaGCN^22^, BayesSpace^23^, stLearn^24^ and spaVAE^25^ modeled the spatial heterogeneity of gene expression to detect these domains. Domains identified by these methods often aligned with anatomical structures visible on histology images accompanying ST data, facilitating the subsequent molecular characterization of these domains^26^. Additionally, novel domains with unique molecular signatures can be detected, frequently leading to new biological discoveries or clinical insights^27^. Unfortunately, while some methods^28,29^, such as BASS and SpatialPCA, could analyze multiple tissue sections, few were explicitly designed to handle spatiotemporal transcriptomics data – spatial transcriptomics datasets collected across multiple time points, which are becoming increasingly popular^28,29^.

Spatiotemporal transcriptomics is an emerging and powerful approach, providing a temporal dimension to characterize the dynamic changes in tissue architecture during development or disease progression. Indeed, spatial domains do not remain static over time. They undergo complex temporal dynamics during development, differentiation, and disease progression, resulting in emergence, disappearance, splitting, and merging of domains over time. For example, during early human brain development, following neural tube closure, three primary domains, known as vesicles -- prosencephalon, mesencephalon, and rhombencephalon -- emerge by embryonic day 28 (E28). By embryonic day 49 (E49), these structures evolve into five secondary brain vesicles, where the prosencephalon divides into the telencephalon and diencephalon, and the rhombencephalon splits into the metencephalon and myelencephalon^30,31^. These changes are important to model and capture as they highlight the dynamic nature of spatial domains as tissues undergo development. However, current computational methods for spatial domain detection are static in nature, ignore temporal information, and fail to model the critical changes that occur during development and disease progression. This gap highlights the critical need for new approaches that can integrate spatial and temporal information to accurately infer spatial domains within tissues and comprehensively characterize the relationship among these domains across time points in spatiotemporal transcriptomics data.

Here, we present SpaDOT (**Spa**tial **DO**main **T**ransition detection), a deep-learning-based computational method designed to accurately identify spatial domains and infer their transitions and relationships across time points in spatiotemporal transcriptomics data. SpaDOT employs a variational autoencoder (VAE) framework integrated with a Gaussian Process prior and graph neighbor information to capture the low-dimensional manifold underlying the data, incorporating hidden clustering variables to characterize distinct spatial domains. In the process, SpaDOT explicitly accounts for both global and local spatial correlation structures among spatial locations within the tissue at each time point and leverages optimal transport (OT) analysis to infer the relationships between spatial domains across time points, thereby enhancing accuracy and interpretation. Through simulation and real data applications, we demonstrate the superiority of SpaDOT in domain detection, latent space preservation, and domain transition tracking over time.

## Results

### SpaDOT Framework

SpaDOT is a computational framework designed for identifying spatial domains in tissues collected at multiple time points and capturing dynamic domain transitions across developmental stages. It takes as input the spatial transcriptomics data along with corresponding spatial coordinates from multiple time points. SpaDOT integrates two complementary approaches -- Gaussian Process (GP) kernel functions and Graph Attention Transformer (GAT) – into a variational autoencoder (VAE) framework to extract spatially aware latent representations, capturing global and local spatial correlation patterns on tissues, respectively (Figure 1, details in Methods). These latent representations are then aligned, harmonized, and constrained using optimal transport (OT) coupling across time points to preserve temporal consistency and infer relationship between domains. Importantly, all modeling steps in SpaDOT are integrated into a unified objective function, ensuring optimal and robust inference of spatial domains across time points and accurate detection of their temporal relationships.

**Figure 1.**
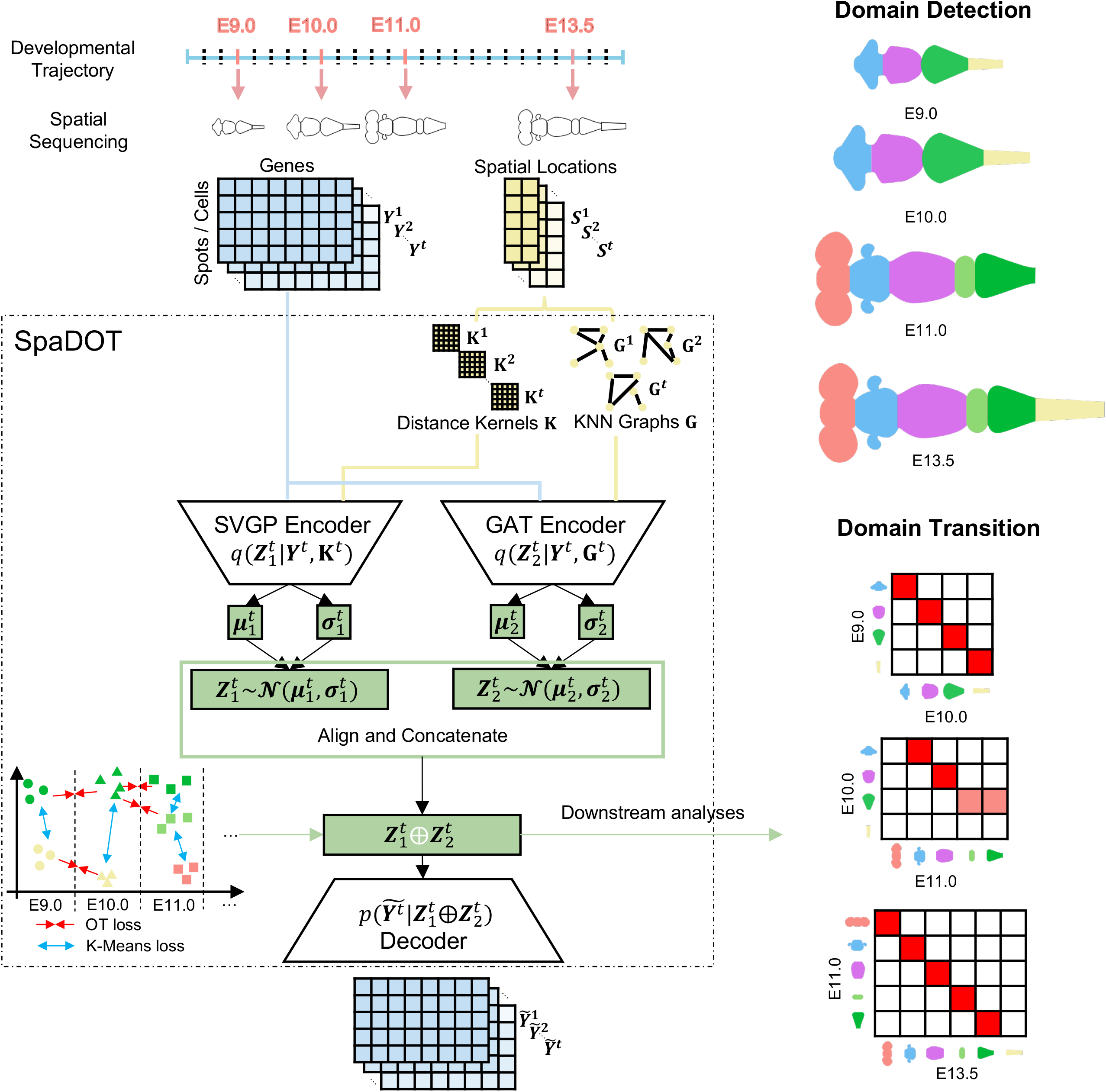
Overview of SpaDOT framework. SpaDOT adopts an integration of two complementary encoders, a Gaussian Process kernel and a Graph Attention Transformer, within one variational autoencoder framework to obtain spatially aware latent representations. The latent representations are further constrained by clustering within each time point and optimal transport (OT) coupling across time points, enabling SpaDOT to identify spatial domains and capture domain transition dynamics. We illustrate SpaDOT’s ability to capture domain splitting, emergence and disappearance along developmental trajectories using the cartoon from brain vesicles development^103^.

### Simulation results

We performed comprehensive simulations to evaluate the performance of SpaDOT (model parameters listed in Supplementary Table S1) and benchmarked it against ten existing methods, which could be categorized into two groups^28^: (1) domain detection methods capable of directly modeling multiple tissue slices, including SpatialPCA^15^, GraphST^32^, SEDR^33^ and spaGCN^22^, and (2) spatial transcriptomics integration methods designed for integrating multiple tissue slices and removing batch effects, including spaVAE^25^, STAligner^34^, PRECAST^35^, spatiAlign^36^, SPIRAL^37^ and STADIA^38^. Since these methods were not designed to model temporal relationships, most could not accommodate varying numbers of spatial domains across time points (Supplementary Table S2), a common scenario in real-world applications of spatiotemporal transcriptomics data. To ensure a fair comparison across methods, we extracted their latent representation and applied K-Means clustering to identify distinct number of domains at each time point.

Simulation details are provided in the Methods section. Briefly, we obtained sagittal mouse brain tissue section from the Allen Mouse Brain Atlas ^39^ at embryonic stage 13.5 (E13.5) and used anatomical annotations to segment the tissue into three, four and five primary domains, modeling the development of an embryonic brain, where three primary domains (forebrain, midbrain and hindbrain) evolve into five domains (telencephalon, diencephalon, midbrain, metencephalon, and myelencephalon) along a temporal development trajectory, with one domain splitting into two at each transition. Particularly, we simulated the forebrain splitting into telencephalon and diencephalon between the first two developmental stages while later on hindbrain splitting into metencephalon and myelencephalon (Figure 2A). In parallel, we segmented single cells from each tissue section using image analysis and simulated their transcriptomic profiles based on a separate scRNA-seq dataset using Splatter^40^. In the simulations, a fraction of genes was differentially expressed (DEGs), either between spatial domains (domain-specific DEGs) or during domain splits (split-induced DEGs), capturing the progression of spatial domains across three time points. We designed four simulation scenarios with varying proportions of domain-specific and split-induced DEGs, each with ten replicates (Figure 2B).

**Figure 2.**
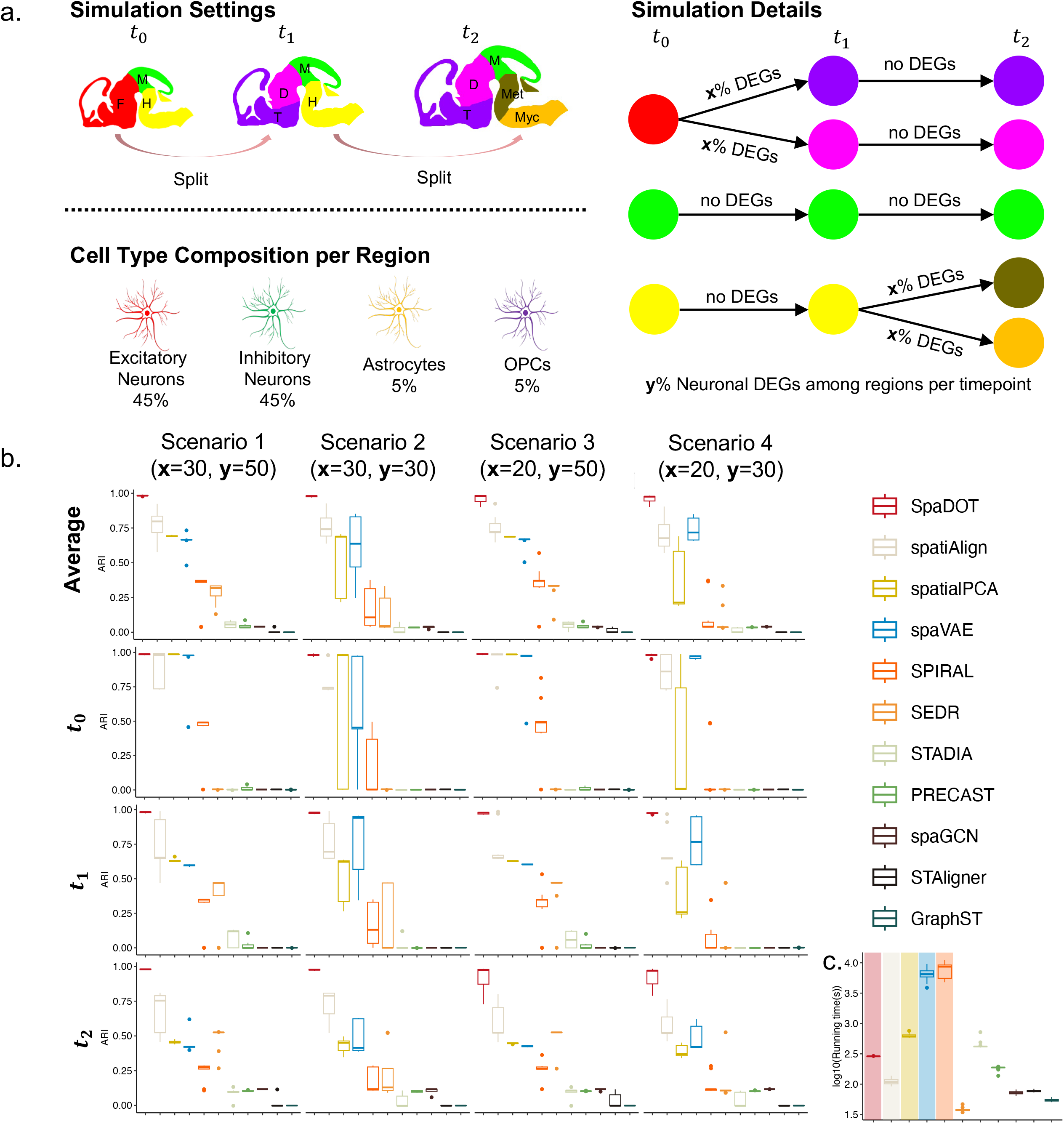
Performance comparisons between SpaDOT and other competing methods on simulated data. **a**. To construct simulations, we obtain sagittal mouse brain tissue from the Allen Mouse Brain at embryonic stage 13.5 (E13.5)^i^ and manually segment it according to annotated biological regions into three, four and five primary domains at time points *t*_0_, *t*_1_ and *t*_2_, respectively. The regions include F – Forebrain, M – Midbrain, H – Hindbrain, T – Telencephalon, D – Diencephalon, Met – Metencephalon, Mye – Myelencephalon. We simulate single-cell transcriptomics data with a fixed cell type composition (45% excitatory neurons, 45% inhibitory neurons, 5% astrocytes and 5% OPCs), and assign cells to each region accordingly. We use **x**% to denote the proportion of split-induced differentially expressed genes (DEGs) and **y**% to denote proportion of domain-specific DEGs. The domain-specific DEGs (**y**%) are only applicable to neuronal cells, following guidance from a mouse whole brain study^69^. We generate four simulation scenarios with different combinations of **x** and **y**, each with ten replicates. **b**. We evaluate domain detection accuracy using adjusted Rand index (ARI) and report ARI-avg across three time points, as well as per time point. Each box in the boxplots represents ten replicates per method. **c**. Execution time (in seconds) for each method is recorded and visualized by aggregating the 40 replicates across the four scenarios regardless of simulation scenarios. The y-axis is log10-transformed. Top five performers in terms of accuracy (ARI) are highlighted. In all boxplots, the center line, box limits and whiskers denote the median, upper and lower quartiles, and 1.5× interquartile range.

We first evaluated spatial domain detection accuracy using three distinct evaluation metrics at each time point: adjusted Rand index (ARI), adjusted mutual information (AMI), and homogeneity (H). Additionally, we calculated the average evaluation metrics across time points, denoted as ARI-avg, AMI-avg, and H-avg. Overall, SpaDOT consistently outperformed the other methods across all scenarios (Figure 2B, Supplementary Figures S1 and S2, Supplementary Table S3). Specifically, SpaDOT achieved a mean ARI-avg of 0.982 and 0.978 (median 0.983 and 0.980) across replicates in the first two easier scenarios, which feature a high proportion of split-induced DEGs. SpaDOT maintained its superior performance in the last two more challenging scenarios, where the proportion of split-induced DEGs was lower, achieving mean ARI-avg of 0.962 and 0.961 (median 0.980 and 0.976; Figure 2B). In comparison, the second-best performer, spatiAlign, achieved mean ARI-avg of 0.771, 0.756, 0.750, and 0.702 (median 0.797, 0.741, 0.722, and 0.677) across the four scenarios, respectively, with similar trends observed in AMI-avg and H-avg. We carefully examined the spatial domains detected by the top three performers -- SpaDOT, spatiAlign, and spaVAE -- along with GraphST, which primarily captured cell types rather than spatial domains, in the most challenging scenario (Scenario 4, Supplementary Figure S3). SpatiAlign identified only the three major domains across time points and failed to capture the domain splits that occur at the two later time points. SpaVAE detected splits at the second time point but missed those between the last two time points. Meanwhile, GraphST clustered cell types but not spatial domains, indicating that spatial information is not fully incorporated^15^. Visualizations of the latent representations inferred by different methods further demonstrate that SpaDOT’s effectiveness in capturing domain dynamics along the temporal trajectory through its latent representations (Supplementary Figure S4). In contrast, the latent representations from spatiAlign and GraphST indicate an over harmonization across time points, obscuring the underlying spatiotemporal dynamics.

Next, we assessed the quality of the detected spatial domains in terms of spatial continuity using three additional metrics at each time point: median local inverse Simpson index (medianLISI), percentage of abnormal spots score (PAS), and spatial CHAOS score (CHAOS). These metrics were again averaged across time points to yield medianLISI-avg, PAS-avg, and CHAOS-avg, respectively. The lower the three scores are, the smoother and more continuous and homogenous domains are. SpaDOT once again achieved the best performance among the methods compared. For example, the mean medianLISI-avg from SpaDOT across ten replicates was 1.000, 1.000, 1.001 and 1.002 (median: 1.000, 1.000, 1.000 and 1.000) for the four scenarios, respectively, indicating a highly homogeneous domain detection. The second best-performer, spatiAlign, achieved a mean medianLISI-avg of 1.079, 1.084, 1.085 and 1.103 (median: 1.083, 1.085, 1.083 and 1.084). Similarly, the mean PAS-avg from SpaDOT across ten replicates was 4.18, 6.82, 2.77 and 7.32 times lower than that of the second best-performer, SpatialPCA, across the four scenarios, respectively. It was also 7.77, 7.57, 5.29 and 6.56 times lower than that of the third best-performer, spatiAlign. Similar trends were observed for CHAOS-avg (Supplementary Figures S5, S6, and S7).

Importantly, a unique feature of SpaDOT is its ability to successfully infer the relationships among the detected domains across time points for all scenarios, accurately capturing the splitting of one domain into two between the two consecutive time points. For example, in a simulation replicate of scenario 1, SpaDOT inferred that the domain 0 (telencephalon) at the second time point originated from domain 0 (forebrain) at the first time point with 0.999 probability, while domain 3 (diencephalon) at the second time point originated from domain 0 (forebrain) at the first time point with probability near 1, supporting the split of domain 0 at the first time point into two domains at the second time point. Similarly, SpaDOT inferred that domain 0 (metencephalon) and domain 4 (myelencephalon) at the third time point originated from domain 1 (hindbrain) at the second time point with probabilities 0.998 and 1, respectively, again capturing the expected domain splits. Similar patterns were observed across the other three scenarios, demonstrating SpaDOT’s ability to consistently and accurately capture domain transitions over time (Supplementary Figure S8). Finally, SpaDOT was proved to be computationally efficient, incurring only a moderate computational cost with an average of 288.204 seconds across scenarios and replicates (Figure 2C). Among the top 5 performers, SpaDOT was only slightly slower than spatiAlign, which had an average computational cost of 110.066 seconds.

### SpaDOT enabled the identification of organogenesis in mouse embryos

We applied SpaDOT to four spatiotemporal transcriptomics studies collected during developmental stages that span diverse species and technologies, including mouse whole embryo and chicken heart measured by 10X Visium^6^, and axolotol telencephalon and mouse whole brain measured by Stereo-seq^41^.

The first dataset was sequenced using 10X Visium and captured spatial transcriptomics measurements during mouse embryonic development at E12.5, E13.5 and E15.5. While the E12.5 and E13.5 slides almost contained the entire embryo, the E15.5 slide primarily covered the head region, with a small portion of heart included. This difference arose due to embryonic growth and tissue expansion at later stages, constrained by the physical size of 10X Visium slides (Figure 3A, Supplementary Figure S9A). As a result, the tissues at the three time points exhibited distinct shapes and morphologies and contained different numbers of spots (2,397, 2,312 and 3,546 at E12.5, E13.5 and E15.5, respectively), posing challenges in identifying spatial domains across time. We applied SpaDOT and other methods to this dataset. Quantitative evaluations demonstrated SpaDOT’s superior performance. Specifically, SpaDOT achieved the highest ARI-avg of 0.586, with ARI values of 0.638, 0.659, and 0.462 for E12.5, E13.5, and E15.5 respectively (Figure 3B, Supplementary Table S4). The second-best performer, SpaGCN, achieved an ARI-avg of 0.530, with ARI values of 0.570, 0.555 and 0.464. SpaDOT also attained the highest AMI-avg of 0.613 (0.684, 0.650 and 0.505) and the highest H-avg of 0.647 (0.693, 0.662 and 0.587), while the second-best performer, spatiAlign, achieved an AMI-avg of 0.576 (0.645, 0.603 and 0.480) and an H-avg of 0.618 (0.662, 0.650 and 0.543). Domain continuity analysis further confirmed the superior performance of SpaDOT (Supplementary Figure S9B). Specifically, SpaDOT achieved a medianLISI-avg of 1.139 (1.239, 1.058, and 1.121), outperforming spaVAE at 1.058 (1.024, 1.106, and 1.044), SPIRAL at 1.108 (1.131, 1.075, and 1.118), and spatiAlign at 1.130 (1.145, 1.169, and 1.076). For PAS and CHAOS, SpaDOT achieved the best PAS-avg of 3.662e-2 (4.172e-2, 2.725e-2, and 4.089e-2), outperforming the second-best spaVAE at 3.772e-2 (3.296e-2, 5.061e-2, and 2.961e-2). Similarly, SpaDOT achieved the lowest CHAOS-avg of 6.276e-2 (6.646e-2, 6.266e-2, and 5.915e-2), with SEDR ranking the second at 6.367e-2 (6.700e-2, 6.380e-2, and 6.021e-2).

**Figure 3.**
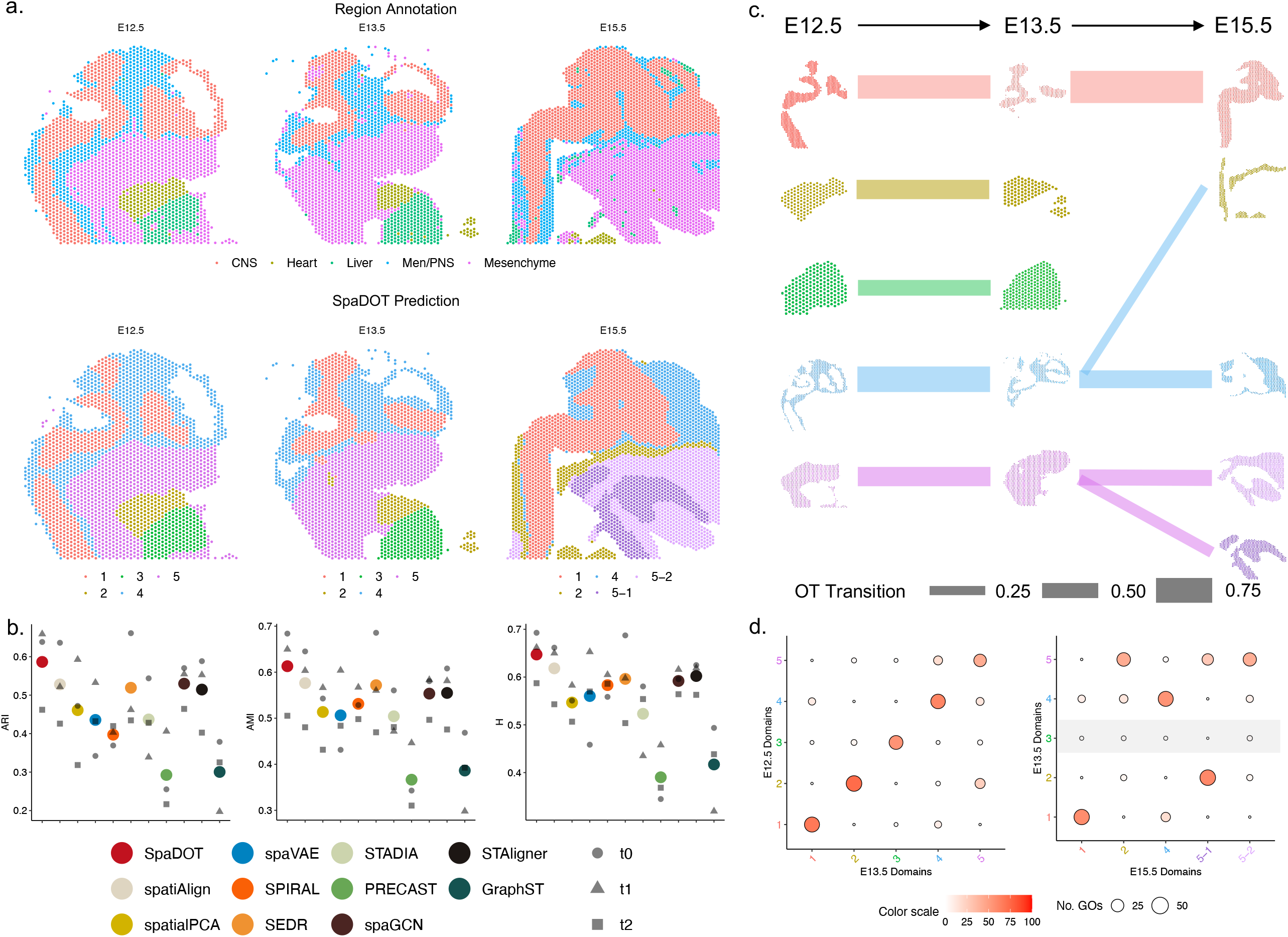
SpaDOT showed mouse organogenesis in mouse embryos. **a**. The original region annotations and spatial domain predictions from SpaDOT at mouse embryonic stage E12.5, E13.5, and E15.5. CNS: Central Nervous System; Men/PNS: Meninges / Peripheral Nervous System. **b**. Domain detection performance evaluated by adjusted Rand index (ARI), adjusted Mutual Information (AMI) and Homogeneity (H). Colored dots represent average values (ARI-avg, AMI-avg and H-avg) across the three time points, while grey dots in different shape correspond to individual performance for each time point. **c**. Optimal transport (OT) transition probabilities between domains across consecutive developmental stages. Transitions are visualized using a flow chart where line widths are scaled by transformation equation: exp(*p*^2^) ∗ 3, where *p* denotes OT transition probability. Transition probabilities less than 0.2 are omitted for clarity. **d**. Dot plot showing the overlap of the top 100 gene ontology (GO) terms enriched in domain marker genes across consecutive time points. Larger and redder the dots indicate a higher number of overlapped GOs. Domain 3 at E13.5 lacks significant GO overlap with any domains at E15.5, suggesting its disappearance. High GO overlap between domains 2 and 5 at E13.5 and domains 2, 5-1, 5-2 at E15.5 indicates functional similarity and shared biological processes.

We carefully examined the domains detected by all methods and observed that SpaDOT was the only method capable of detecting the heart, liver and surrounding mesenchymal region at E12.5 and E13.5, where the three organs were present on the tissue, while the other methods failed (Figure 3A, Supplementary Figure S10A). For example, while SEDR and SpatialPCA partially delineated the boundaries between heart and liver at E12.5, both methods either fused the two organs into one single domain or missed one domain entirely at E13.5. The effective detection of the three domains was attributed to SpaDOT’s integration of two complementary latent representation frameworks, Gaussian Process (GP) kernel function and Graph Attention Transformer (GAT), as the former captured smoother spatial patterns across both time points while the later enhanced the detection of sharper domain boundaries despite its occasional overlook on the finer details necessary for resolving specific regions (Supplementary Figure S10B). At E15.5, SpaDOT captured the major anatomical regions in the brain and distinguished the mesenchymal region into two distinct domains, one of which was also consistently detected by most methods except for spatiAlign and spaVAE. The detection of these two mesenchymal subregions was supported by the hematoxylin and eosin (H&E) stained image (Supplementary Figure S9A), which accurately delineated the domain boundaries detected by SpaDOT. Additionally, differential gene expression and functional enrichment analyses further validated these two subregions, with one significantly enriched in muscle system-related biological processes and the other in skeletal system-related processes (Supplementary Figure S11).

Beyond accurately detecting domains across time points, another key advantage of SpaDOT is its ability to leverage optimal transport (OT) transition probabilities to capture the relationships among domains over time (Figure 3C, Supplementary Figure S12A, transition probabilities calculation details in Methods section). For example, SpaDOT identified domain 4 at E15.5 as primarily derived from domain 4 at E13.5, with a transition probability of 0.584, which in turn predominantly originated from domain 4 at E12.5, with a transition probability of 0.765. Functional enrichment analysis revealed those domains labeled as 4 from E12.5, E13.5, and E15.5 shared enrichment in gene ontology (GO) terms related to forebrain development, telencephalon development, pallium development, and neural precursor cell proliferation (Supplementary Figure S12B). Similarly, SpaDOT detected that domain 1 at E15.5 primarily derived from domain 1 at E13.5, which in turn originated from domain 1 at E12.5, with transition probabilities of 0.881 and 0.721, respectively. These domains were enriched in biological processes related to axonogenesis, cell projection organization, and microtubule cytoskeleton regulation, all of which are associated with neuronal development and connectivity (Supplementary Figure S11C). The number of overlapping GO terms across these clusters were statistically significant: 131 overlapping terms (permutation test Z-score=4.558, p-value = 5.17e-6) and 141 overlapping terms (permutation test Z-score=9.076, p-value < 2.2e-16), indicating a consistency of domain detection (Supplementary Figure S11D, permutation test details in Methods section). Importantly, through GO analysis, we observed a notable increase in the number of time point-specific GO terms at E15.5 domains, indicating increasing complexity and specialization at this later developmental stage. Specifically, domain 4 in E15.5 contained 282 time point-specific GO terms, compared to 62 in E13.5 and 87 in E13.5. Similarly, domain 1 in E15.5 included 143 unique GO terms, exceeding 18 in E12.5 and 25 in E13.5. For example, domain marker genes from domain 4 at E15.5 were enriched in GO terms related to cerebral cortex development and neuron-specific processes, such as regulation of neuron differentiation, cerebral cortex neuron differentiation, and neuron migration. These findings suggest an active period for cortical layer development and highlight the emergency of more specialized biological functions at E15.5. Supporting evidence from the literature confirmed that most callosal projection neurons (CPN) were generated between E14.5 and E16.5, with these late-born CPN migrating to superficial cortical layers during this specific time period^42,43^.

In addition to capturing domain transition over time, SpaDOT effectively detected the expected domain disappearance from E13.5 to E15.5. Specifically, the transition probabilities for domain 3 (liver) at E13.5 to domains detected at E15.5 were low, indicating the disappearance of the liver at E15.5 (Figure 3C). Furthermore, GO enrichment analysis revealed few overlaps of enriched GO terms between liver at E13.5 and other domains detected at E15.5, confirming the liver’s disappearance (Figure 3D, details in Method section). Both observations are aligned with the fact that the E15.5 slide does not include the liver region. Additionally, SpaDOT did not identify a separate heart domain at E15.5 due to its extremely small size (37 out of 3,546 spots, 1.043%), instead merging it into the surrounding mesenchymal tissue (domain 2). This result was supported by the original annotation, gene expression profile dissimilarity, and minimal GO term overlaps between this mesenchymal subregion (domain 2) and another two subregions (domains 5-1 and 5-2; Supplementary Figure S12E-F). Indeed, domain 2 at E15.5 exhibited 110 unique GO terms, including processes related to the transforming growth factor-beta (TGF-β) signaling pathway and endothelial cell proliferation, suggesting a potential role in vascular development^44^.

### SpaDOT revealed valvulogenesis during chicken heart cardiogenesis

We further applied SpaDOT to a spatiotemporal transcriptomics dataset focusing on the heart region of chicken embryos profiled using 10X Visium^45^. Unlike mouse embryos, chicken hearts are smaller, allowing multiple replicates to be analyzed on a single slide, providing an opportunity for comparative analysis. This dataset measured four key Hamburger-Hamilton (HH) ventricular development stages: day 4 (HH21-HH24), day 7 (HH30-HH31), day 10 (HH35-HH36) and day 14 (∼HH40). On day 4, the early chamber emerged with the initiation of ventricular septation, primarily consisting of trabeculated myocardium. By day 7, the four-chamber heart formed with more defined ventricular septation and fenestrated trabeculated sheets. On day 10, the septation was complete, and myocardium became the dominant tissue component. By day 14, cardiogenesis concluded, resulting in a fully developed four-chamber heart with compact myocardium^45^.

We applied SpaDOT along with other methods to this dataset, obtaining tissue domains for each developmental stage (Figure 4A, Supplementary Figure S13, Supplementary Table S5). SpaDOT consistently demonstrated superior performance across all metrics (Figure 4B, Supplementary Figure S14). In terms of domain detection accuracy, SpaDOT ranked first with an ARI-avg of 0.530, achieving ARI values of 0.444, 0.383, 0.615, and 0.677 on days 4, 7, 10, and 14, respectively. In comparison, STADIA ranked the second with an ARI-avg of 0.505, attaining ARI values of 0.459, 0.443, 0.464, and 0.652. SpaDOT achieved the second-best AMI-avg at 0.573 (0.450, 0.534, 0.673 and 0.636), slightly below STADIA’s 0.582 (0.504, 0.592, 0.595 and 0.639), while attaining the highest H-avg of 0.571 (0.455, 0.538, 0.672 and 0.619), compared to STADIA’s 0.541 (0.490, 0.564, 0.523 and 0.589). Despite STADIA’s seemingly competitive performance on these metrics, further inspection revealed its limitations: it completely missed domain 4 on both day 7 and day 10, and failed to identify valves, which SpaDOT correctly identified as domains 5 on both day 7 and day 10 (Figure 4A, Supplementary Figure S13). Additionally, the domains detected by SpaDOT were spatially more continuous than STADIA: SpaDOT achieved a medianLISI-avg of 1.647 (1.851, 1.942, 1.564 and 1.232), PAS-avg of 0.141 (0.131, 0.210, 0.112 and 0.110), and CHAOS-avg of 7.489e-2 (7.844e-2, 7.079e-2, 6.610e-2 and 8.424e-2); while STADIA attained a medianLISI-avg of 1.733 (2.288, 1.895, 1.571 and 1.177), PAS-avg of 0.235 (0.402, 0.255, 0.157 and 0.127, ranking 7th), and CHAOS-avg of 7.791e-2 (8.397e-2, 7.271e-2, 6.946e-2 and 8.549e-2, ranking 7th).

**Figure 4.**
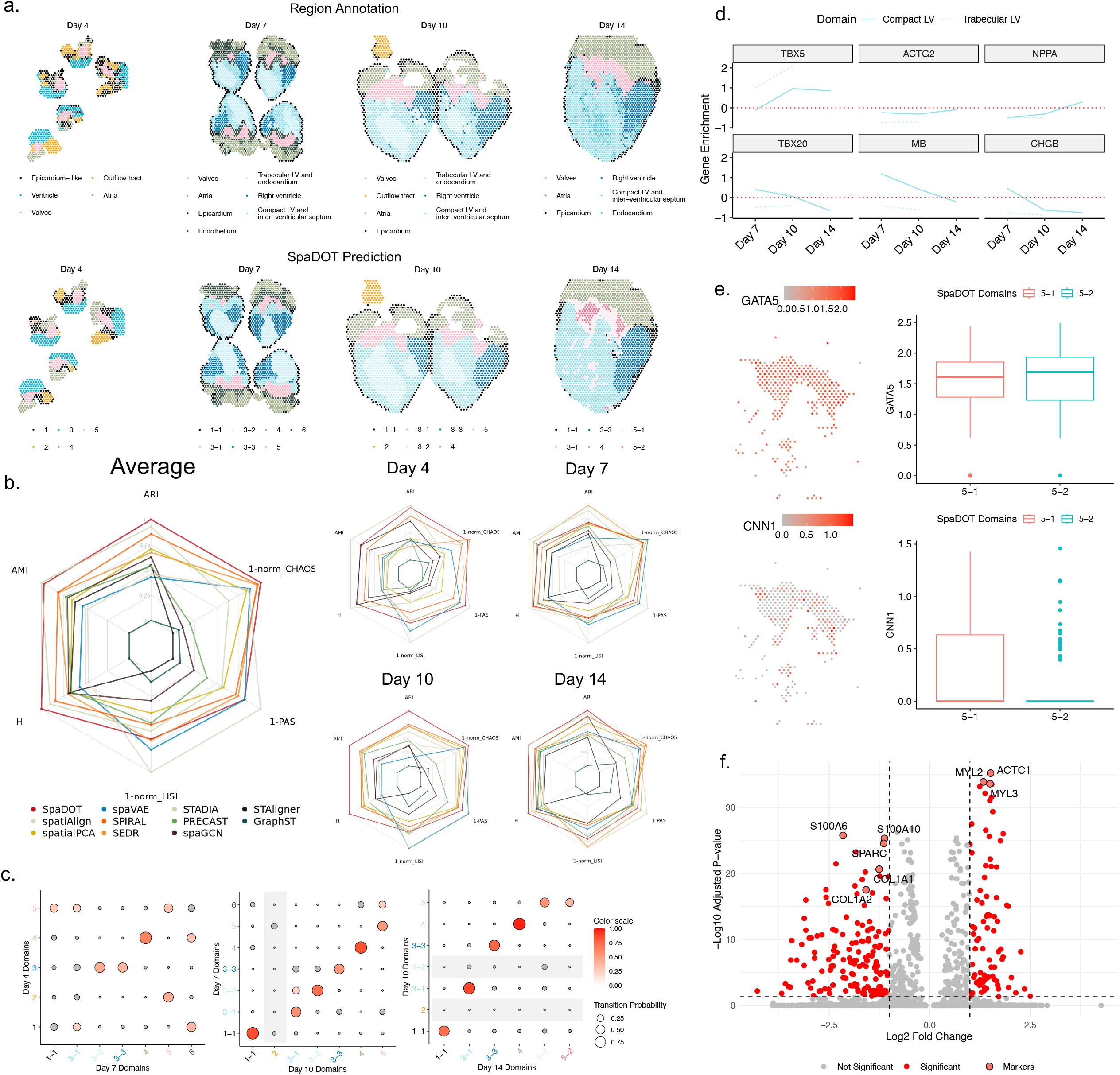
SpaDOT identified valvulogenesis in developing chicken heart. **a**. The original region annotations and spatial domain predictions from SpaDOT across four developmental time points Day 4, 7, 10 and 14 in chicken heart. **b**. Radar plots summarize domain detection performance in terms of accuracy and spatial continuity. Accuracy metrics include adjusted Rand index (ARI), adjusted Mutual Information (AMI), Homogeneity (H); Spatial continuity metrics include 1-normalized median local inverse Simpson index (1-norm_LISI), 1 – percentage of abnormal spots score (PAS) and 1 - normalized spatial CHAOS score (1 – norm_CHAOS). Both average scores across time points and individual time points results are presented. **c**. Dot plots of the optimal transport (OT) transition probabilities between domains across consecutive time points. Rows or columns with no transition probabilities > 0.2 are shaded in grey, indicating either the emergence of new domains or the disappearance of existing domains. For example, domain 2 emerges at Day 10 and domains 2 and 3-2 disappear by Day 14. **d**. Gene enrichment analysis comparing compact left ventricle (Compact LV) and trabecular left ventricle (Trabecular LV), using the right ventricle as a baseline. Due to the disappearance of Trabecular LV, gene expression enrichment trends are only shown for Day 7 and Day 10 for Trabecular LV. **e**. Marker gene expression across valvular domains 5-1 and 5-2. *GATA5* is a general valve marker gene, while *CNN1* is a marker gene specific to atrioventricular valves. Boxplots show the gene expression distributions of *GATA5* and *CNN1* across domains. In all boxplots, the center line, box limits and whiskers denote the median, upper and lower quartiles, and 1.5× interquartile range. **f**. Volcano plots display the differentially expressed genes (DEGs) between domains 5-1 and 5-2 at Day 14. DEGs are considered as significant using an adjusted p-value threshold of 0.05 and an absolute log2 Fold Change > 1. Upregulated genes in domain 5-1 appear on the right, while upregulated genes in domain 5-2 appear on the left. Top three DEGs for each domain along with key collagen fiber genes that indicate the identity of semilunar valves are highlighted.

We carefully assessed the domain dynamics inferred by SpaDOT during development and found that it successfully identified several expected dynamic patterns across different ventricles, including the right ventricle, left compact ventricle, and left trabecular ventricle, along the time trajectory (Figure 4C). Specifically, on day 7, the right ventricle (domain 3-3) exhibited a high transition probability of 0.545 to domain 3-3 on day 10, and subsequently, a high transition probability of 0.753 to domain 3-3 to day 14. The compact left ventricle (domain 3-1) on day 7 showed a high transition probability of 0.562 to domain 3-1 on day 10, followed by 0.876 to domain 3-1 on day 14. The trabecular left ventricle (domain 3-2) on day 7 exhibited a high transition probability of 0.703 to domain 3-2 on day 10. As development progresses, the trabecular ventricle largely disappeared and transited into the endocardium.^45,46^. We conducted the gene enrichment analysis of the two left ventricles compared to the right ventricle using marker genes from the literature^45^ (Figure 4D) and found that the gene expression trend confirmed our observations: chromogranin B (*CHGB*) was highly enriched in the right ventricle from day 7 onwards, and T-box transcription factor 5 (*TBX5*) showed enrichment in the left ventricle, with a decrease from day 7 to day 14. These trends further support the accuracy of the domain detection by SpaDOT in the ventricles. Additionally, GO enrichment analysis on the right ventricle, left compact ventricle, and trabecular ventricle showed highly similar top enriched GO terms (Supplementary Figure S15).

SpaDOT identified several important domain emergence and disappearance events during development. First, the outflow tract, absent from the tissue on day 7, was captured on the tissue section on day 10, but disappeared again on day 14 due to the limited tissue coverage of 10X Visium. The transition matrices from SpaDOT effectively captured this progression: a new domain (domain 2) emerged on day 10, with no significant transition probabilities from any domains detected on day 7, and disappeared on day 14, with its domains showing low transition probabilities from domain 2 on day 10. This captured dynamic highlights SpaDOT’s ability to accurately track domain transitions along temporal trajectories. Second, SpaDOT accurately captured a domain split event between day 10 and day 14, where the valve (domain 5) on day 10 differentiates into two distinct domains (domains 5-1 and 5-2) on day 14. This reflects the valvulogenesis^47^, the development of two valve types, atrioventricular valves (include tricuspid and mitral valves) and semilunar valves (include aortic and pulmonary valves)^46,48–50^: the two types of valve begin to distinguish from each other structurally around day 5.5 (HH28), with semilunar valves fully formed by stage HH34 and atrioventricular valves fully formed by day 10 (HH36), with atrioventricular valves experiencing some additional remodeling in later stages between day 10 and day 14 (HH36-40). To validate this, we first checked the valve marker gene *GATA5* provided in the original study^45^ and confirmed that it was highly enriched in both domains on day 14, supporting their identity of being valves (Figure 4E). Then, we found that a atrioventricular valve-specific marker gene, *CNN1*^51^, was highly expressed in domain 5-1, while the mitral valve marker gene, *CRABP2*, an atrioventricular valve subtype specific marker, was not expressed. This suggests that domain 5-1 likely corresponds to tricuspid valve, an atrioventricular valve subtype (Figure 4E). Differential expression analysis between domains 5-1 and 5-2 confirmed the cardiomyocytes identity of the tricuspid valve^52,53^ (domain 5-1; Figure 4F; Supplementary Figure S16A). Supporting this, a study on the chicken heart has found that, unlike the mitral valve, the tricuspid band receives contribution from both endocardial cushion tissue and ventricular myocardium, which contain endocardial-derived cells and cardiomyocytes^54,55^. Additionally, *ATP2A2*, which is homologous to *Serca2* in mouse^49,56^, has been shown to express in some β-galactosidase-positive cells within the atrioventricular leaflets, but not in semilunar valves (Supplementary Figure S16B). In contrast to atrioventricular valves, semilunar valves are more fibrous that mainly contain collagen fibers type I (*COL1A1* and *COL1A2*) and type III (*COL3A1*). Visualization of these genes showed their significant enrichment in domain 5-2 compared to domain 5-1, which, along with their high ranking among differentially expressed genes (*COL1A1* ranks 6^th^ and *COL1A2* ranks 13^th^; Figure 4F), supported the identification of domain 5-2 being semilunar valves (Figure 4F; Supplementary Figure S16B)^57,58^.

### SpaDOT captured neurogenesis in axolotl telencephalon

We next applied SpaDOT to a Stereo-seq data from the axolotl telencephalon across three developmental stages (Stages 44, 54, and 57)^41^, capturing key transitions during the development of the pallium into the medial pallium (MP), dorsal pallium (DP), and lateral pallium (LP; Figure 5A). Compared to 10X Visium, Stereo-seq offers a larger chip size (ranging from 1 cm × 1 cm to 13 cm × 13 cm), which allows for the capture of full tissue anatomy to ensure enhanced spatial domain consistency across all three stages^59^ (Figure 5A). We applied SpaDOT alongside other methods to identify spatial domains (Supplementary Figure S17, Supplementary Table S6). For domain detection accuracy, SpaDOT outperformed all other methods, achieving the highest ARI-avg of 0.649, with ARI scores of 0.715, 0.627, and 0.606 at consecutive time points. It also ranked the first in AMI-avg (0.667; 0.664, 0.678, and 0.660) and H-avg (0.678; 0.659, 0.694, and 0.682). The second-best method in ARI was SEDR, with an ARI-avg of 0.566 (0.607, 0.574, and 0.517). For AMI-avg and H-avg, SPIRAL ranked the second, achieving an AMI-avg of 0.626 (0.584, 0.647, and 0.649) and an H-avg of 0.649 (0.600, 0.666, and 0.680) (Figure 5B). SpaDOT detected continuous spatial domains, with a medianLISI-avg of 1.331 (1.322, 1.360, and 1.311), comparable to SEDR (1.274; 1.294, 1.339, and 1.189). SpaDOT also achieved the lowest CHAOS-avg of 5.482e-2 (7.514e-2, 5.179e-2, and 3.754e-2), whereas the second-best method, spatiAlign, attained a CHAOS-avg of 5.567e-2 (7.635e-2, 5.278e-2, and 3.788e-2). Additionally, SpaDOT achieved the lowest PAS-avg of 0.056 (0.050, 0.061, and 0.059), with spaVAE ranking second at 0.065 (0.081, 0.058, and 0.054) (Supplementary Figure S18A). Notably, SpaDOT exhibited the least variance in medianLISI and PAS scores across the three time points, highlighting its stable performance over time.

**Figure 5.**
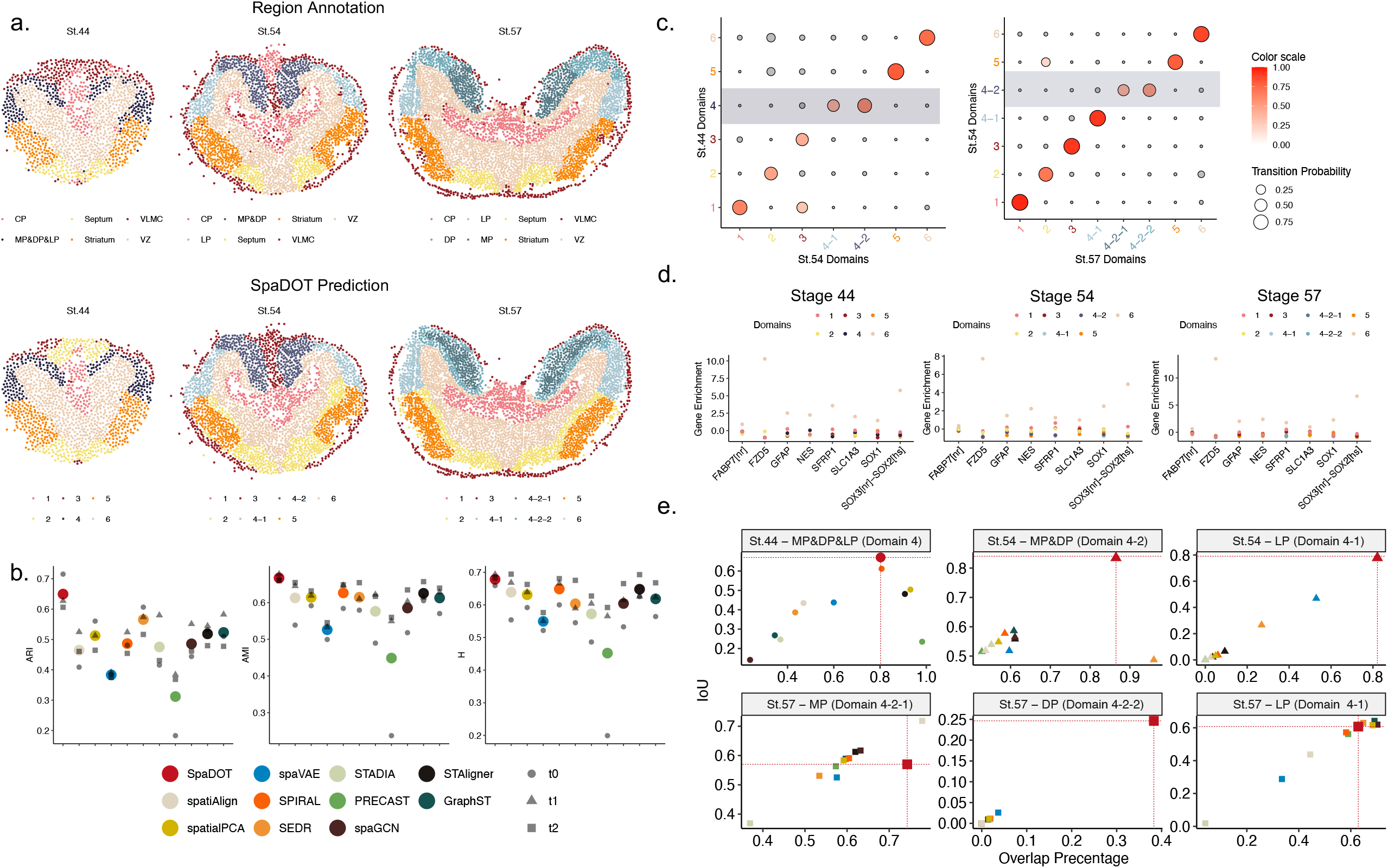
SpaDOT captured domain evolution during Axolotl telencephalon neurogenesis. **a**. The original region annotations and spatial domain predictions from SpaDOT across three developmental stages 44, 54 and 57 (St.44, St.54 and St.57) in axolotl telencephalon. **b**. Domain detection performance evaluated by adjusted Rand index (ARI), adjusted Mutual Information (AMI) and Homogeneity (H). Colored dots represent average values (ARI-avg, AMI-avg and H-avg) across the three time points, while grey dots in different shape correspond to individual performance for each time point. **c**. Dot plots of the optimal transport (OT) transition probabilities between domains across consecutive time points. Shaded area outline domain splits, such as domain 4 at St.44 splits into domains 4-1 and 4-2 at St.54 and domain 4-2 at St.54 splits into domains 4-2-1 and 4-2-2. **d**. Gene enrichment analysis of ventricular zone (VZ) markers across time points. The consistent high enrichment in domain 6 at all stages suggest domain specificity related to VZ. **e**. Scatter plots comparing the overlap percentage and Intersection over Union (IoU) between predicted domains and annotated regions. The dotted lines outline SpaDOT’s performance; methods appearing further to the right and higher up demonstrate better domain detection. Ideally, a method consistently shows the top-right corner across domains within each individual time point indicates robust and accurate detection.

Importantly, SpaDOT stood out as the only method capable of preserving the integrity of the ventricular zone (VZ; domain 6) across all three time points (Supplementary Figure S17). In contrast, other methods primarily detected various cell types within the VZ instead of identifying the complete region as a standalone domain. Moreover, SpaDOT ensured temporal continuity of the detected VZ, with a high transition probability of 0.731 from Stage 44 to Stage 54, followed by a high transition probability of 0.874 from Stage 54 to Stage 57 (Figure 5C). We conducted differential gene expression analysis to examine the transcriptomic signatures of the detected VZ at different stages and identified *Slc1a3, Gfap*, and *Sox2* as the top enriched genes at each developmental stage. *Slc1a3* and *Sox2* have been found as established markers for neural stem cells, while *Gfap* serves as an astroglia marker, all of which play crucial roles in neurogenesis during development^60–62^. Subsequent marker gene enrichment analysis and GO analysis further confirmed the importance of neurogenesis, with mitotic cell cycle phase transition terms along with DNA replication-related terms being highly enriched across all stages, indicating rapid proliferation and differentiation of neuronal stem cells (Figure 5D, Supplementary Figure S18B). The VZ domains at different stages also shared a considerable overlap of GO terms, supporting their functional similarity (74 out of 332; permutation test Z-score = 2.428, p-value = 0.015; Supplementary Figure S18C).

SpaDOT also represented the only method that successfully identified pallium split, which was the domain split from MP&DP&LP (domain 4) at Stage 44 to LP (domain 4-1) and MP&DP (domain 4-2) at Stage 54 (Figure 5C). Specifically, SpaDOT detected a high transition probability of 0.377 (to domain 4-1) and 0.574 (to domain 4-2) from domain 4 at Stage 44. Additionally, SpaDOT provided evidence, albeit weak, on another expected split from MP&DP (domain 4-2) at Stage 54 to MP (domain 4-2-1) and DP (domain 4-2-2) at Stage 57. We used Hungarian algorithm^63^ to create a one-to-one mapping between the predicted domain labels and annotations based on the overlap percentage for each method (calculated as the number of overlapping spots divided by the total number of spots in the predicted domain). We then computed the Intersection over Union (IoU) with the mapped domain labels for each method and centered on MP, DP, LP related domains. The results showed that SpaDOT outperformed the other methods, achieving both the highest IoU and the greatest overlap percentage for Stages 44 and 54 (Figure 5E). For Stage 57, SpaDOT was the only method that identified distinct domains for MP, DP and LP. Although spatiAlign performed better than SpaDOT in detecting MP, it failed to distinguish DP and LP as separate domains. Similarly, SEDR, SpatialPCA, SpaGCN and GraphST all failed to separate MP and DP. To understand why MP, DP and LP were hard to be distinguished, we calculated the gene expression similarity of those pallium regions across time points at Stage 57. We noticed that even in the original annotation, related regions across time points were not clustered together (Supplementary Figure S19A; details in Methods section). While in SpaDOT, we noticed a high similarity among domain 4 (MP&DP&LP) at Stage 44, domain 4-2 (MP&DP) at Stage 54, and domain 4-2-2 (DP) at Stage 57 (Supplementary Figure S19B). Spearman correlations between these domains were high: 0.893 between domain 4 at Stage 44 and domain 4-2 at Stage 54, 0.908 between domain 4 at Stage 44 and domain 4-2-2 at Stage 57, and 0.895 between domain 4-2 at Stage 54 and domain 4-2-2 at Stage 57. In the meantime, domain 4-1 at Stage 54 (LP) and Stage 57 (LP) were clustered together with a high Spearman correlation of 0.904, indicating a stabilized and specific LP expression during the developmental stage. Compared to all clusters, domain 4-2-1 (MP) at Stage 57 only had a high Spearman correlation with domain 4-2-2 at Stage 57 (0.886), suggesting a unique function of this specific region. We further investigated the potential function of domain 4-2-1 by performing differential gene expression analysis between domain 4-2-1 and domain 4-2-2. Notably, we observed that *CBLN2*, a known member of the retinoic acid (RA) signaling pathway, was highly enriched in domain 4-2-1 (Supplementary Figure S19C)^64^. Additional GO analysis and enrichR^52^ analysis under the CellMarker 2024^53^ category consistently highlighted synaptic activities, further supporting that domain 4-2-1 might play a unique role in embryonic development and neurogenesis.

### SpaDOT detected the dynamic change of thalamus domain during mouse brain development

We applied SpaDOT to a developing mouse embryonic dataset from Stereo-seq, measuring mouse whole embryos collected from E9.5 to E16.5 with embryonic stage 1 as interval ^65^. The anatomic annotations were available at two levels of granularities: organ-level (e.g. brain, kidney, heart) and cell-type level (e.g. mucosal epithelium and olfactory epithelium, which contains specific epithelial cell types^66^). Due to the different granularity levels of the annotation, we focused our analysis on one organ – the brain – and examined brain development between E12.5 and E16.5. Since the number of spatial domains at each time point was unknown, which was common in most real-world scenarios, we estimated the number of spatial domains for each time point using the elbow method and applied these values to all methods for comparison (details in Methods; Supplementary Figure S20A). This yielded optimal domain numbers of 7, 8, 10, 9 and 10 for the five time points in SpaDOT, respectively (Figure 6A, Supplementary Figure S21). To obtain the identity of predicted domains, we used marker genes from the original study^8^ to visualize expression patterns and calculated enrichment within each detected domain across time points (Supplementary Figure S20B, S20C). Based on the marker genes, we manually curated the region identities. Overall, the detected regions by SpaDOT exhibited relatively exclusive prediction assignments and with higher marker gene enrichment (Figure 6B) compared to SpatialPCA, especially in the first two stages (Supplementary Figures S20C, S22A).

**Figure 6.**
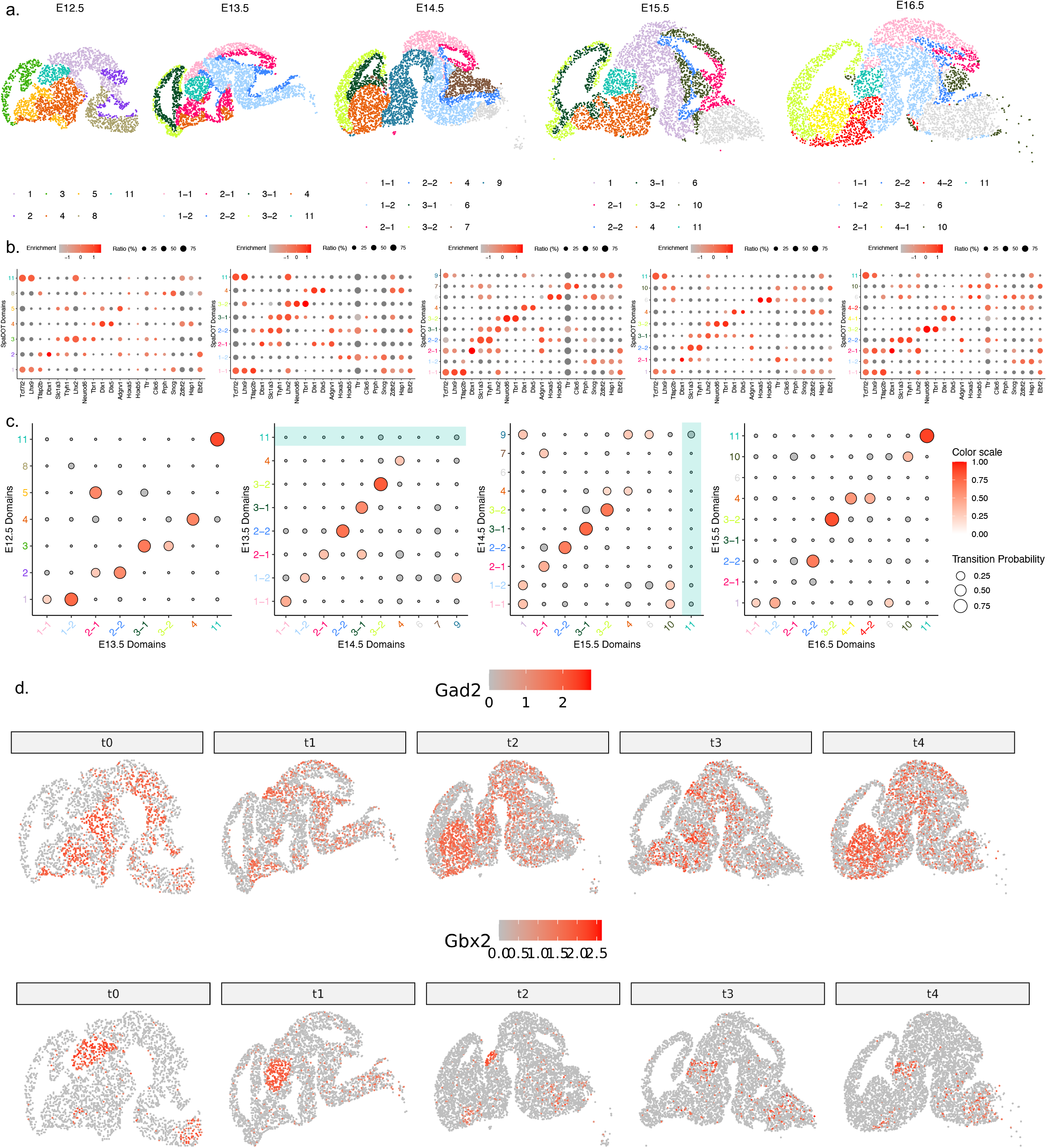
SpaDOT revealed domain emergence and disappearance during mouse brain development. **a**. The spatial domain predictions from SpaDOT across five mouse embryonic stages in mouse brain, including E12.5, E13.5, E14.5, E15.5 and E16.5. **b**. Dot plots show marker gene enrichment for each domain predicted by SpaDOT. Dot color intensity reflects enrichment levels and dot size indicates the proportion of spots within the domain expressing the marker genes. **c**. Dot plots of the optimal transport (OT) transition probabilities between domains across consecutive time points. Shaded areas outline the disappearance of the thalamus at E14.5 and its re-emergence at E15.5 (domain 11), likely due to segmentation slice location. **d**. Spatial expression patterns of key marker genes across all embryonic stages. *Gad2* is a GABAergic neuron marker and *Gbx* is a thalamus marker.

In terms of spatial continuity, SpaDOT ranked the highest across all three metrics (Figure 6B, Supplementary Table S7). Specifically, SpaDOT achieved the best medianLISI-avg of 1.080, with LISI values of 1.048, 1.182, 1.056, 1.078 and 1.034 across embryonic stages, outperforming SPIRAL (1.080; 1.079, 1.161, 1.168, 1.070 and 1.082) and SpatialPCA (1.100; 1.111, 1.197, 1.070. 1.054 and 1.070). For PAS, SpaDOT achieved an average of 3.831e-2 (2.541e-2, 4.752e-2, 4.153e-2, 3.705e-2 and 4.002e-2), while SpatialPCA ranked the second with an average of 4.668e-2 (6.145e-2, 7.306e-2, 3.337e-2, 2.810e-2 and 3.744e-2), and SPIRAL ranked the third with an average of 8.132e-2 (5.583e-2, 8.241e-2, 9.078e-2, 9.438e-2 and 8.322e-2). For CHAOS, SpaDOT again achieved the best performance with an average value of 3.072e-2 (3.649e-2, 3.210e-2, 2.854e-2, 2.786e-2 and 2.862e-2) followed by SpatialPCA (CHAOS-avg: 3.099e-2, 3.782e-2, 3.274e-2, 2.813e-2, 2.769e-2 and 2.858e-2) and GraphST (CHAOS-avg: 3.248e-2, 3.753e-2, 3.250e-2, 3.144e-2, 2.961e-2 and 3.132e-2). Although SpatialPCA showed the closest performances to SpaDOT for all three metrics, the domains detected by SpatialPCA along with marker gene enrichment analysis (Supplementary Figures S21 and S22) indicated inferior performance. SpatialPCA failed to detect pallium domain at E13.5 and combined pallium ventricular zone and pallium together as domain 2. Furthermore, at E16.5, it incorrectly separated pallium into two domains (domain 4 and domain 10), which was inconsistent with the pallium marker genes expression patterns provided in the paper^65^ (*Neurod6* and *Tbr1*; Supplementary Figure S23A).

SpaDOT achieved consistent domain identification across time points and detected domain appearance, disappearance and reappearance events through OT analysis. For example, a unique region - domain 11, which was consistently detected at E12.5, E13.5, E15.5, and E16.5, disappeared at E14.5. These regions showed a lack of expression of Glutamate decarboxylase 2 (*Gad2*), a GABAergic neuron marker, while exhibiting enrichment of *Gbx2* and *Rora*, both known as thalamus markers (Figure 6D, Supplementary Figure S24) ^67,68^. Additionally, findings from a scRNA-seq mouse whole brain atlas confirmed the predominance of glutamatergic neurons in the thalamus^69^. The enrichment of *Lhx9*, a marker from a human thalamus study, further supported the identification of these regions at all time points except E14.5 (Supplementary Figure S24) ^70^. The detection of the appearance and disappearance of the thalamus further supported the assumption that certain tissue regions might not be fully captured due to technical artifacts in some segmentation sections. As a result, it is essential to incorporate flexibility into domain detection methods developed for spatial transcriptomics data, rather than assuming a fixed number of domains in the model.

## Discussion

In this paper, we introduce SpaDOT, a model designed to analyze spatiotemporal transcriptomics datasets and provide insights into spatial domain development along temporal trajectories. SpaDOT uniquely leverages spatial location information using two complementary approaches: a Gaussian Process (GP) kernel function for capturing global, smooth patterns and a Graph Attention Transformer (GAT) for detecting local, discrete patterns. By integrating both within a variational autoencoder (VAE) framework, SpaDOT facilitates flexible, end-to-end latent embedding learning. Unlike existing multi-slice integration methods that do not account for temporal information^28^, SpaDOT demonstrates superior domain detection results across diverse datasets spanning different sequencing technologies (10X Visium and Stereo-seq), species (mouse, axolotl, and chicken), and tissue types (brain, telencephalon, heart, and whole embryo). Notably, SpaDOT is the only method that explicitly accounts for temporal dynamics using optimal transport (OT), while accommodating varying numbers of spatial domains at each time point - an important feature not supported by other methods, such as STAMP^71^. Additionally, in contrast to other multi-slices integration models that tend to over-harmonize slices, SpaDOT preserves the unique characteristics of individual time points while accurately identifying domain transitions. Furthermore, SpaDOT represents a novel advancement by focusing on inferring domain-level developmental trajectories rather than spot- or cell-level changes. This sets it apart from existing approaches like stLearn^24^, SpaceFlow^72^ and SPATA2^73^, which are designed for spot-level trajectory analysis to infer pseudo-spatiotemporal patterns within a single ST tissue slice. By modeling domain-level temporal transitions, including splitting, emergence, and disappearance, SpaDOT adopts a broader perspective and effectively captures key developmental events, as revealed by the transition probability matrices derived from its OT component.

There are several possible extensions to SpaDOT. First, SpaDOT is not only accurate in detecting spatial domains and highly effective in revealing their dynamics over time, but also computationally efficient, completing analyses for moderate-sized datasets within minutes. However, as spatiotemporal transcriptomics studies continue to grow in size, further improvements in computational efficiency will enhance the scalability and usability of SpaDOT. The primary computational bottleneck arises from the incorporation of GP prior, which in its naïve form incurs a cubic time complexity of *O*(*n*^3^), where *n* is the number of spots^74^. To address this, we have implemented a stochastic variational Gaussian Process (SVGP) ^25,75^ that uses inducing points to reduce the time complexity to *O*(*nm*^2^), where *m* is the number of inducing points, which can be set proportionally across time points to ensure that larger slices receive more inducing points. If necessary, additional computational gains could be achieved by introducing sparse kernels (e.g., set a cutoff distance) or by applying low-rank approximations to the kernels, as implemented in SpatialPCA^15^. Beyond improving the GP component, replacing the Graph Attention Transformer (GAT) with more computationally efficient alternatives could further enhance SpaDOT’s scalability. Promising alternatives include the simplified graph convolutional network^76^, the weighted light graph convolution network^77^, and the graph convolutional network via initial residual and identity mapping layers^78^, all of which use fewer parameters and can substantially reduce training time. Second, determining the number of spatial domains remains a common challenge in the field^14,79^. During the training stage, because the true number of domains is unknown, SpaDOT uses a K-Means loss with a preset large number of domains. This strategy generates well-separated latent representations and avoids the need to predefine the number of domains. Empirically, we find that the K-Means loss is effective in encouraging the formation of distinct clusters. During the inference stage, when the number of domains is unknown, we apply a combination of the elbow method and the within-cluster sum of squares (WSS) metric, using an arbitrary threshold to identify the optimal number. Certainly, K-Means may not always yield the most optimal clustering result^80^. Community-based clustering methods, such as Louvain and Leiden^81,82^, also allow users to specify the number of domains and may outperform K-Means on certain datasets. For consistency, we have used K-Means in SpaDOT during training across all datasets analyzed in the present study. However, the SpaDOT framework is flexible and can readily accommodate alternative clustering methods. Users are encouraged to explore alternative clustering methods on the latent representations learned from SpaDOT to achieve optimal performance. Finally, SpaDOT represents a pioneering effort in identifying spatial domain dynamics along with temporal trajectories in spatiotemporal transcriptomics datasets. As such, it is not intended for application to spatial transcriptomics (ST) data collected at a single time point and is best suited for datasets that include multiple time points to ensure robust identification of functional domains. By incorporating clustering and time constraints, SpaDOT detects both shared domains across time points as well as domain transitions between consecutive time points. In summary, SpaDOT provides a much-needed entry point for exploring how spatial domains evolve over time, providing a strong foundation for future research into dynamic spatial transcriptomic processes.

## Methods

### SpaDOT overview

SpaDOT was specifically designed for spatial transcriptomics studies to identify spatial domains in tissues and uncover their dynamic progression across multiple time points, representing different developmental stages or stages of disease progression. SpaDOT accounted for multiple dynamic processes, including the splitting of existing spatial domains, the emergence of new domains, the disappearance of certain domains, and other complex transitions that would occur in various biological settings over time. This capability enabled spaDOT to comprehensively capture and characterize intricate changes in tissue architecture throughout these stages. To achieve this, spaDOT first projected the high-dimensional spatial omics data into a low-dimensional manifold by synergizing two complementary encoders within variational autoencoders (VAE) framework: (1) a conditional deep probabilistic encoder conditioned on spatial coordinates, which captured global spatial patterns and encourages spatial smoothness across tissue locations; (2) a graph-structured deep probabilistic encoder, which captured neighborhood interactions and discrete spatial patterns. Together, these two encoders extracted complementary information from the spatial omics data, enabling effective manifold encoding. For the conditional probabilistic encoder, we adopted a Gaussian Process (GP)-based VAE model, namely the stochastic variational Gaussian Process Variational Autoencoder (SVGPVAE)^75^, where spatial correlation was incorporated into the covariance function of the GP. For the graph-based probabilistic encoder, we adopted Graph Attention Variational Autoencoder (GATVAE) that operated on a K-Nearest Neighbor (KNN) graph constructed from spatial coordinates, where K was set to a small number (e.g. 6) to focus on capturing local structures. We harmonized the latent variables derived from the two encoders by aligning their scales and concatenating them into a unified latent space, which was further passed through a shared decoder. On this unified low-dimensional manifold, we applied two additional constraints: a clustering constraint based on the K-Means loss function, which partitioned the tissue into well-separated potentially functional spatial domains, and a temporal constraint based on the optimal transport (OT) loss function, which established temporal couplings between spatial domains across consecutive time points, thus allowing SpaDOT to capture the dynamic progression of functional domains while maintaining spatial and temporal consistency. Importantly, the embedding process through the encoders and the application of constraints were jointly integrated into a single loss function, ensuring effective optimization and seamless coordination between the two steps. SpaDOT was also designed as an end-to-end trainable framework, ensuring scalability to accommodate the growing volumes of spatiotemporal transcriptomics datasets.

### SpaDOT details

We assumed that spatial transcriptomics data were collected on the same tissue at *T* different time points. At each time point *t* (*t* ∈ {1, ⋯, *T*}), the spatial transcriptomics data ***Y***^*t*^ consisted of a gene expression matrix with dimensions *m*^*t*^ × *n*^*t*^, where *m*^*t*^ denoted the number of genes, and *n*^*t*^ denoted the number of measured spatial locations. We used feature selection (details in the next section - data preprocessing and gene selection) to derive a common set of *g* genes shared across all time points, therefore 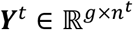. In addition to the gene expression matrix, the spatial coordinates for each data were represented by ***S***^*t*^, with dimensions 2 × *n*^*t*^. Utilizing the input ***Y***^*t*^ and ***S***^*t*^, our goal was to first obtain an interpretable and accurate low-dimensional latent space denoted as ***Z***^*t*^, with dimensions *d* × *n*^*t*^, where *d* was shared among time points. With the learned latent space, we further inferred functional domains represented as ***R***^*t*^, with dimension *r*^*t*^ × *n*^*t*^, at each time point. To achieve both goals, SpaDOT utilized three distinct loss functions:

- Latent Loss denoted by **L**_Latent_
- K-Means Loss denoted by **L**_KMeans_
- OT Loss denoted by **L**_OT_

The first loss function, **L**_Latent_, trained an integrated deep probabilistic model to derive effective latent space for the data at each time point, represented as ***Z***^*t*^. The second loss function, **L**_KMeans_, inferred a relatively large number of candidate spatial domains (e.g., *r*^*t*^ = 10 at time point *t*), ***R***^*t*^, from the latent space ***Z***^*t*^ at each time point through K-Means clustering. In the meantime, the third loss function, **L**_OT_, encouraged similarity in the latent space representations of the same spatial domain across consecutive time points, ensuring temporal consistency in the inferred spatial domains.

### Loss **L**_Latent_

Building upon the foundation of SVGPVAE and GATVAE, this loss function trained a generative model capable of generating ***Y***^*t*^ given ***S***^*t*^, while concurrently obtaining an interpretable low-dimensional latent space ***Z***^*t*^ that can be later employed for functional domain detection. Each 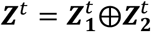 was a concatenation of the latent space obtained from SVGPVAE and GATVAE, denoted as 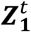 and 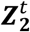 respectively. For simplification, we focused on only one time point and dropped the upper script time point *t* for illustration purpose.

When reconstructing gene expression profiles, we adopted a shared decoder that took the aligned and concatenated latent space as input and we defined one reconstruction loss 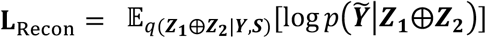 to quantify the differences between the reconstructed data 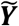 and the original data ***Y***. We employed the Mean Squared Error (MSE) as the reconstruction metric.

In SVGPVAE, we introduced a conditional prior, *p*(***Z***_**1**_|***S***), and computed the Kullback-Leibler (KL) divergence between the approximated posterior and the prior. The divergence was denoted as **L**_SVGPVAE-KL_ = D_KL_ (*q*(***Z***_**1**_|***Y***) ∥ *p*(***Z***_**1**_|***S***)). We modeled the *p*(***Z***_**1**_|***S***) with GP prior by positing *d*_1_ independent latent functions with 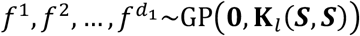 with a kernel function parameterized by spatial locations, and therefore 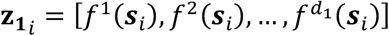. The 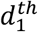 latent channel of all latent variables 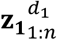 followed covariance **K**_*l*_ (***S, S***). Here, we selected radial basis function (RBF) kernel 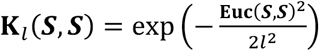 where **Euc**(·,·) indicated *l* Euclidean distance, and *l* was the length scale of the kernel. We set *d*_^1^_ = 10 and *l* = 0.1 by default. When dealing with a large amount of data, calculating the exact GP posterior may encounter computational burden. To ensure the model’s scalability, a sparse variational GP^83^ was employed to approximate the full GP by choosing a set of inducing points and the approximation was fulfilled by variational inference. More details in the Supplementary Note 1.

In GATVAE, we construct a KNN undirected graph as input, denoted by **G** = (***V, E***), where ***V*** represents a set of vertices and ***E*** represents a set of edges connecting the nodes. Each vertex is a spatial spot / cell, and each edge indicates whether the spatial spots are within the closest *k* neighbors based on spatial distance. We set the number of neighbors to be 6 ∗ round(*n*/1000) for each time point. We represent the graph as an adjacency matrix ***A*** where ***A***_*ij*_ = 1 indicates an edge between vertex *i* and vertex *j* and ***A***_*ij*_ = 0 otherwise. The graph attention encoder dynamically aggregates features from local neighborhoods, assigning adaptive attention scores to each neighbor. The approximated posterior in GATVAE is denoted by *q*(***Z***_**2**_|***Y, A***) and we assume the prior *p*(***Z***_**2**_) follows a standard multivariate Gaussian distribution ***Z***_**2**_**∼**𝒩(**0, I**). We then compute the KL divergence as **L**_GATVAE-KL_ = D_KL_ (*q*(***Z***_**2**_|***Y, A***) ∥ *p*(***Z***_**2**_)). We set the size *d*_2_ of latent space ***Z***_**2**_ to 10. More details can be found in Supplementary Note 1.

The latent spaces ***Z***_**1**_ and ***Z***_**2**_ generated by two different encoders were expected to exhibit discrepancies in scale due to differences in their architectures and parameters. These scale mismatches can hinder direct comparisons and latent space alignments. To address this, we introduced an alignment loss **L**_Alignment_ to harmonize the latent spaces. We computed the *ℓ*_2_-norm of ***Z***_**1**_ and ***Z***_**2**_ along the columns that resulted in two vectors of size *n*. Each *ℓ*_2_-norm was normalized by the number of dimensions *d*_^1^_ and *d*_2_, respectively. The alignment loss **L**_Alignment_ was then defined as the MSE between these two vectors to ensure consistent scaling:

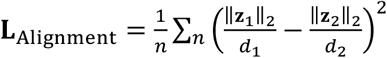. In the end, we obtained a latent space with dimension (*d*_^1^_ + *d*_2_) × *n*, where we set *d*_^1^_ = 10 and *d*_2_ = 10. In summary, the latent loss was expressed as **L**_Latent_ = **L**_Recon_ + *β*_1_**L**_SVGPVAE-KL_ + *β*_2_**L**_GATVAE-KL_ + *ω*_1_**L**_Alignment_ where *β*_1_, *β*_2_ and *ω*_1_ were weighting factors.

#### Loss **L**_KMeans_

To obtain spatial domain information, we applied K-Means clustering on the pre-trained embedding space ***Z***^*t*^ following the K-Means cost function demonstrated in IRIS^10^ to link the embeddings ***Z***^*t*^ to spatial domains ***R***^*t*^. Different from IRIS, our achievement of introducing K-Means was twofold, one was to obtain distinct spatial domains, and the other was to stabilize the latent space across training epochs within the same time point. Since the exact number of clusters were unknown during the training stage, we set K=10 and applied K-Means to latent space ***Z***^*t*^ derived at each time point to obtain 10 cluster centroids 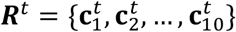 where each 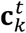 represented the mean of the data points assigned to the *k*-th cluster and the dimension was *d* × 1. Based on the K-Means clustering results, we introduced a hidden binary indicator variable indicating cluster assignment 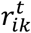. If a spot *i* was assigned to domain *k*,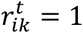. otherwise, 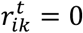. The centroids were computed as 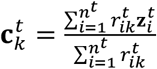 and the K-Means loss was computed as 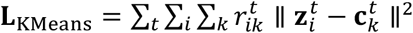.

#### Loss **L**_OT_

Besides considering embeddings within the same time point, we introduced unbalanced optimal transport (OT)^84,85^ to track the dynamic change of spatial domains on the tissue along the temporal trajectory across multiple time points. Based on the K-Means clustering centroids 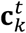 obtained at time point *t* and centroids 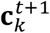 obtained at the next following time point *t* + 1, we computed a transport plan **P**^*t,t*+1^, with dimension *k*^*t*^ × *k*^*t*+1^ (where *k*^*t*^, *k*^*t*+1^ were set as 10), containing a set of couplings 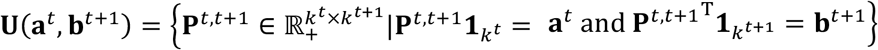. Here, the row sums of **P**^*t*,t+1^ equaled to the source distribution **a**^*t*^ at time point and the column sums of **P**^*t,t*+1^ equaled to the target distribution **b**^*t*+1^. Due to the domain development across time points, **a**^*t*^ and **b**^*t*+1^ might have different total masses, we therefore introduced entropy regularizations and unbalanced OT regularizations to penalize the loss function for calculating **P**^*t,t*+1^:

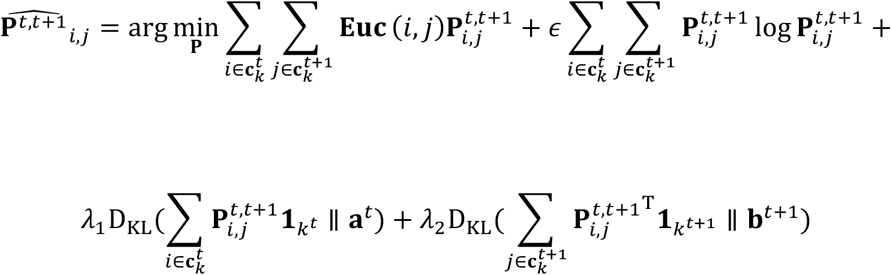

We set *λ*_1_ = 0.1, *λ*_2_ = 10 and *ϵ* = 0.01 to promote row imbalance while ensuring a relatively confirmative transition. With the calculated transport plan 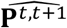, the loss function of OT became 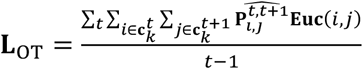. The usage of cluster-wise OT loss enhanced computational efficiency and allowed for more efficient domain tracking across time points.

The final loss function of SpaDOT was a sum of the above three terms: **L**_SpaDOT_ = **L**_Latent_ + *ω*_2_**L**_KMeans_ + *ω*_3_**L**_OT_ = **L**_Recon_ + *β*_1_**L**_SVGPVAE-KL_ + *β*_2_**L**_GATVAE-KL_ + *ω*_1_**L**_Alignment_ + *ω*_2_**L**_KMeans_ + *ω*_3_**L**_OT_. Based on the scale of each loss component, we set default weighting parameters as follows: *β*_1_ = 1, *β*_2_ = 1*e* − 4, *ω*_1_ = 0.1, *ω*_2_ = 0.1 and *ω*_3_ = 1. The hyperparameters ensured that the losses were comparable in magnitude, providing a balanced contribution to the optimization. For user convenience, these hyperparameters were set default but can be optionally modified.

### Data preprocessing and gene selection

For the input spatial gene expression count matrix, we first performed library size normalization to a total number of 10,000^86^ and performed log1p transformation separately for each time point. We then selected spatially variable genes (SVGs) using SPARK-X^87^ with default parameters and used adjusted p-value less than 0.05 as cutoff to select significant SVGs. The number of SVGs detected for each time point was denoted as 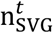. Based on the selected SVGs, we first performed PCA to obtain the top 30 PCs and then built a graph based on KNN using 100 as perplexity. We used Louvain clustering algorithm with resolution as 1.0 on the KNN graph to obtain community structure. If the number of clusters 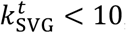, we increased the resolution by 0.1 until 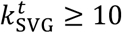. When 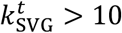, we merged the detected clusters using *mergeCommunities* function from *bcluster* R package^88^ to let the number of clusters equal to 10. We then kept 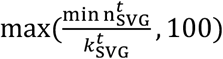 number of SVGs within each detected clusters and unioned those SVGs across time points as feature genes. Based on selected feature genes, we performed center scale transformation. As for the spatial coordinates, we performed standardization by removing the mean and scaling to unit variance. The center scaled data, time point indicator, and standardized spatial coordinates were used as input for SpaDOT.

### Simulation study

We conducted simulation study to evaluate the performance of our method. Background. To mimic a real biological situation, we simulated the process of brain development where three primary brain regions gave rises to five secondary brain regions during embryonic development^31,89^. Originally, the three primary brain regions included the prosencephalon (forebrain), mesencephalon (midbrain), and rhombencephalon (hindbrain). Later on, the forebrain was divided into two parts: telencephalon and diencephalon, where the telencephalon later formed the cerebral hemispheres, and the diencephalon later formed structures like thalamus and hypothalamus. The hindbrain gave rise to two parts as well: metencephalon and myelencephalon, where the metencephalon later formed the pons and cerebellum and the myelencephalon later formed the medulla oblongata.

#### Simulation details on constructing spatiotemporal tissue images

Based on the prior background knowledge, we designed a three-timepoint functional region split (Figure 2A). The construction process followed the simulation procedure demonstrated in SpatialPCA^15^ with modifications. We first extracted a reference tissue contour with Adobe Photoshop from a developing mouse brain collected at E13.5 stage from Allen Reference Atlas – Developing Mouse Brain^39^ (https://atlas.brain-map.org/atlas?atlas=181276130). Through the Interactive Atlas Viewer, we clearly identified three main brain regions: forebrain (F), midbrain (M) and hindbrain (H). With this information, we manually segmented the tissue image into three spatial domains and set this segmented image as the reference image for the first timepoint (*t*_0_). For the second timepoint (*t*_1_), we assumed that the forebrain gave rise to two parts as indicated in the Interactive Atlas Viewer as secondary prosencephalon (SP) and diencephalon (D). Again, we manually segmented the forebrain region (F) into two separate regions which resulted in four spatial domains at *t*_1_. Similarly, for the third timepoint (*t*_2_), we assumed that hindbrain gave rise to two parts where one contained the prepontine hindbrain (PPH) and pontine hindbrain (PH) while the other one contained the pontomedullary hindbrain (PMH) and medullary hindbrain (MH). After manually segmenting the hindbrain region (H) into two separate regions, we obtained five spatial domains at *t*_2_. To mimic the growth of brain, based on the same image canvas size, we set the smallest occupation proportion for *t*_0_, a larger proportion for *t*_1_ and the largest proportion for *t*_2_. We randomly sampled 9,000, 12,000 and 15,000 pixels from images at timepoints *t*_0_, *t*_1_ and *t*_2_ to serve as single-cell locations and obtained their spatial coordinates respectively.

#### Simulation details on constructing corresponding gene expressions

With the single-cell locations along with their coordinates, we aimed to simulate gene expression for each location. We took a scRNA-seq study on human prefrontal cortex obtained via Smartseq2^90^ as reference and we used *Splatter* R package^91^ to estimate the cell-type-specific parameters for Neurons, GABAergic neurons, Astrocytes and Oligodendrocytes Precursor Cells (OPCs). We assumed that each spatial region contained the same cell type composition that consists of the above four cell types: Neurons (45%), GABAergic neurons (45%), Astrocytes (5%) and OPCs (5%). According to a recent study of whole mouse brain^92^, neuronal cells have been reported to hold the characteristics of being region-specific while non-neuronal cells shared most features among brain regions. Therefore, we considered a certain amount of differentially expressed genes (DEGs) among spatial domains from the same timepoint for neuronal cells while no DEGs for non-neuronal cells. In *Splatter*, this was controlled by parameters *de*.*prob* and for different simulation scenarios, we set *de*.*prob* to be 0.3 and 0.5 to indicate the difference among spatial domains (domain-specific DEGs) within the same timepoint. As for the domain split across timepoint (e.g. forebrain splits into telencephalon and diencephalon; denoted as F splits into SP and D in anatomy), we considered it as batch effect for all four cell types and we used *batch*.*facLoc* from *Splatter* to control for it. For different simulation scenarios, we set *batch*.*facLoc* to be 0.2 and 0.3 (split-induced DEGs). Together, we obtained a combination of four simulation scenarios and for each scenario, we obtained a set of gene expression profiles for each cell type at each timepoint. We then randomly assigned the simulated gene expression profiles to each spatial locations on the image based on the cell type composition (45% for Neurons, 45% for GABAergic neurons, 5% for Astrocytes and 5% for OPCs) and generated the single-cell resolution spatial transcriptomics data. In total, we simulated 100,000 cells with 5,000 genes including 45,000 Neurons, 45,000 GABAergic neurons, 10,000 Astrocytes and 10,000 OPCs. For a faster simulation result, we down sampled the single-cell spatial locations to 2,250, 3,000, and 3,750 locations and randomly assigned cells to each location according to the abovementioned cell type composition.

### Real data analyses

We applied SpaDOT analysis on four spatiotemporal transcriptomics studies, namely Mouse Organogenesis Dataset, Chick Cardiogenesis Dataset, Axolotl Telencephalon Neurogenesis Dataset, and Mouse Brain Neurogenesis Dataset. We manually adjusted the domain labels and colors of SpaDOT for better visualization purpose.

#### The Mouse Organogenesis Dataset^93^

This dataset discovered the regulatory programs during mouse organogenesis by measuring spatial gene expression via 10X Visium and spatial ATAC-seq. We focused on the spatial gene expression where the dataset collected mouse embryo at three embryonic days: E12.5, E13.5 and E15.5, with 2397, 2312 and 3546 spots respectively. Each time point contained one slide / replicate and shared 22,496 genes. After performing feature selection procedure, we had 9,481 genes remained.

When performing marker gene enrichment for Central Nervous System (CNS) and Men/PNS (Meningeal / Peripheral Nervous System), we noticed that certain genes had extremely high enrichment (Fold change > 30) in heart region such as *Cox8b, Nppb, Itk* and *Myh7*. We therefore removed them in our analyses accordingly. When calculating domain-specific marker genes, we considered a gene as marker gene using the cutoff of adjusted p-values less than 0.05 and average log2 fold change larger than 1. Then in the Gene Ontology (GO) enrichment analysis, we selected top 100 genes per domain. For GO terms overlapping analysis, we selected top 100 enriched GO terms and performed the overlap analysis for domains between consecutive time points.

#### The Chicken Cardiogenesis Dataset^45^

This dataset measured chicken heart development from early to late four-chambered heart stage by measuring spatial transcriptomics by 10X Visium. In total, there were four time points collected, including Day 4 (HH21-HH24), Day 7 (HH30-HH31), Day 10 (HH35-HH36) and Day 14 (∼HH40), where each time point corresponded to one slide. However, due to the slide size, Day 4 contained 5 replicates, Day 7 contained 4 replicates, Day 10 contained 2 replicates and Day 14 only contained 1 replicate. There were 747, 1966, 1916 and 1967 spots at each time point and the number of genes measured were 23,986 genes. After performing feature selection procedure, we had 2,771 genes remained.

When calculating domain-specific marker genes, we considered a gene as marker gene using the cutoff of adjusted p-values less than 0.05 and average log2 fold change larger than 0. We used all marker genes to perform GO enrichment analysis for each domain. When visualizing the radar plot, since medianLISIs were larger than 1, we first performed 0-1 normalization across methods to make the minimum value of LISI to be 0 and maximum value of LISI to be 1. Then we used 1 minus the normalized LISI to compute 1-norm_LISI for each time point and then computed the average. We conducted the same procedure for CHAOS. For PAS, since they were normalized between 0 and 1, we directly used 1 minus the original score.

#### The Axolotl Telencephalon Neurogenesis Dataset^41^

This dataset discovered cellular and molecular features of the axolotl telencephalon during development and injured-regeneration by measuring spatial transcriptomics by Stereo-seq. In our study, we only focused on developmental stages and due to the earlier split of MP, DP and LP regions, we subset the whole developmental dataset to three stages: stage 44, stage 54 and stage 57, where each stage contained 1477, 2929 and 4410 spots. In the original study, MP and DP were incorrectly annotated as separate spatial domains at Stage 44 before their actual separation, which began at Stage 54. To better reflect the developmental process, we redefined the domains by merging MP, DP, and LP into a single domain (MP&DP&LP) at Stage 44 and combining MP and DP into one domain (MP&DP) at Stage 54. This adjustment captured the sequential domain splitting: a single domain at Stage 44 divided into MP&DP and LP at Stage 54, with MP&DP further separating into MP and DP by Stage 57. In the end, each stage contained one slide / replicate and shared 12,704 genes. After performing feature selection procedure, 2,198 genes remained.

When calculating domain-specific marker genes, we considered a gene as a marker gene using the cutoff of adjusted p-values less than 0.05 and average log2 fold change larger than 1. Since the dataset lacks gene annotations for axolotls, we converted gene symbols to mouse gene symbol prior to conducting GO analysis. We selected top 50 genes per domain to perform GO analysis.

#### The Mouse Whole Brain Dataset^8^

The dataset was a subset of mouse MOSTA dataset. The original data contained the mouse whole embryo from time point E9.5 to E16.5 and each had multiple technical replicates. We focused on embryonic stages between E12.5 and E16.5 with one replicate at each time point, including E12.5 (E1S2), E13.5 (E1S2), E14.5 (E1S1), E15.5 (E1S1) and E16.5 (E1S1). Then, we kept those spots assigned as *Brain* in the original annotation and removed certain outliers with the spatial coordinates *y* (E12.5: no filter, E13.5: *y* < -450, E14.5: *y* < -500, E15.5: *y* < 200, E16.5: *y* < -575). In the end, we obtained 73,916 spots and 26,580 genes. For a quicker result, we down sampled 20,000 spots and after down sampling, there were 2597, 3325, 4729, 4588 and 4698 spots at each embryonic stage respectively and after performing feature selection, 7,622 genes remained.

When determining the number of domains for each time point, we used the elbow method on within-cluster sum of squares (WSS) as metric, which was a compatible metric to the K-Means loss. We computed WSS for K-Means clusters from 5 clusters to 20 clusters and then computed the WSS difference between consecutive **K**. We then computed the ratio between the consecutive difference to detect the most gradient change to detect the elbow point. With the slope getting slower, in case we always detect a minor elbow point, we set a threshold of having WSS difference to be larger than 20 and then we selected the number of clusters with highest ratio. In the end, we obtained 7 (WSS diff:73.029, ratio: 2.189), 8 (WSS diff: 64.407, ratio: 2.136), 10 (WSS diff: 41.829, ratio: 1.771), 9 (WSS diff: 26.084, ratio: 2.230), 10 (WSS diff: 38.159, ratio: 1.669) for E12.5, E13.5, E14.5, E15.5 and E16.5 respectively.

### Benchmark analyses

We benchmarked SpaDOT to other competing methods that are capable of performing multi-slices integration of spatial transcriptomics datasets on both simulation study and the real data analysis. Some of the methods were designed for spatial clustering with multi-slices clustering option such as: SpatialPCA^15^ (version 1.3.0, https://github.com/shangll123/SpatialPCA), GraphST^94^ (version 1.1.1, https://github.com/JinmiaoChenLab/GraphST), SEDR^33^ (version 1.0.0, https://github.com/JinmiaoChenLab/SEDR), and spaGCN^22^ (version 1.2.7, https://github.com/jianhuupenn/SpaGCN). Other methods explicitly accounted for batch effect among multi-slices including spaVAE^25^ (https://github.com/ttgump/spaVAE), STAligner^95^, PRECAST^96^ (version 1.6.5, https://github.com/feiyoung/PRECAST), spatiAlign^97^ (version 1.0.0, https://github.com/zhoux85/STAligner), SPIRAL^98^ (version 1.0, https://github.com/guott15/SPIRAL) and STADIA^99^ (version 1.0.1, https://github.com/yanfang-li/stadia). Following a recent benchmark study^100^, we evaluated the performance from three perspectives: (1) spatial domain identification along with functional analysis; (2) continuity analysis of spatial domains; and (3) domain transition analysis along time trajectory.

#### Spatial domain identification

For a fair comparison on domain identification, we obtained embeddings from these methods and performed K-Means clustering based on the number of given annotated clusters within each time point. We used Adjusted Rand Index (ARI), Normalized Mutual Information (NMI), Adjusted Mutual Information (AMI) and Homogeneity (H) to evaluate the clustering performance within each time point. We also averaged these values across time points for each metric to generate ARI-avg, NMI-avg, AMI-avg and H-avg as an overall metric for comparison.

#### Continuity analysis of spatial domains

The continuity analysis of spatial domains aims to measure the spatial coherence of spatial domains and whether the domains share clear boundaries. Following SpatialPCA^15^, we utilized predicted spatial domains along with spatial coordinates to evaluate the continuity. Median local inverse Simpson index (medianLISI), CHAOS, and percentage of abnormal spots (PAS) were used as evaluation metrics for each tissue slice at each time point. For all three metrics, smaller values indicate better spatial continuity and better spot homogeneity within spatial cluster as well as more homogeneous neighborhood spatial domain clusters of the spot. We averaged these values across time points for each metric to generate medianLISI-avg, CHAOS-avg, and PAS-avg.

#### Domain transition analysis along time trajectory

With the latent space ***Z***^*t*^ and the predicted domains ***R***^*t*^ at each time point *t*, we computed a spot-wise optimal transport (OT) plan matrix **P**^(*t,t*+1),Spot)^ between consecutive time points *t* and *t* + 1. This matrix was computed under the imbalance constraints as described in Waddington-OT, with parameters *λ*_1_ = 1, *λ*_2_ = 50 and *ϵ* = 0.01. The reason the parameters are larger than in **L**_OT_ is because a larger constraint can help with eliminate noisy signals when conducting spot-level OT analysis. The weaker constraint on the row imbalance compared to column imbalance reflects biological assumption that spatial spots from time point *t* might disappear in the next following time point *t* + 1. In the computed matrix **P**^(*t,t*+1),Spot)^, each row represented the distribution of mass transported from a specific spot at time point *t* to all spots at time point *t* + 1, while each column represented the distribution of mass received by a specific spot at time point *t* + 1 from all spots at time point *t*. Then, we aggregated the **P**^(*t,t*+1),Spot)^ into **P**^(*t,t*+1),Domain)^ of dimensions *k*^*t*^ × *k*^*t*+1^ according to the predicted spatial domain assignments for each spot. For spots residing in each domain 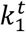, we first computed the total mass transported to the next time point *t* + 1 by 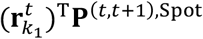 where 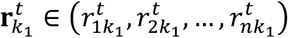 and 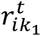 is a binary indicator variable indicating whether spot *i* belongs to cluster 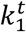. Then for each domain 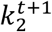 at time point *t* + 1, we had a vector 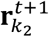. We then derived the domain-wise OT plan matrix 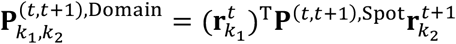. Finally, we performed two normalizations on **P**^(*t,t*+1),^ by normalizing each row sum to 1 and each column sum to 1, interpreting the entries as transition probabilities. We calculated the 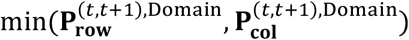 of the two normalized matrices to obtain the final transition probability matrix. When interpreting the matrix, each column represented the probability distribution of a target domain at time point *t* + 1 originating from the source domains at time point *t*, providing a probabilistic interpretation of the developmental trajectories between domains across consecutive time points. We visualized the transition probabilities using heatmaps and dot plots and we considered those transition probabilities less than 0.2 as noises. When interpreting from a row perspective: if a spatial domain remains unchanged between time points, its corresponding row in the heat map will display a single high transition probability, indicating a direct mapping. When a domain splits into multiple domains or a new domain emerges from an existing domain, the corresponding row will display two or more high transition probabilities. In contrast, if a domain disappears, its row will not exhibit any high transition probabilities, indicating a lack of contribution to any target domains. When interpreting from a column perspective: if values from a column do not show high transition probabilities, the indication is a new domain emerges.

#### Biological downstream analysis

In addition to computational comparisons, we validated the detected spatial domains through downstream analysis. Specifically, for each time point, we identified marker genes for each predicted domain using the *FindAllMarkers* function from the *Seurat* R package (v5.1.0)^101^. For Gene Ontology (GO) enrichment analysis, we utilized the *enrichGO* function from *clusterProfiler* R package (v4.8.3)^102^ with selected domain-specific marker genes from each domain. We use the *BP* as the subontology, *BH* for adjusting p-value and a q-value cutoff of 0.05 for all analysis. The corresponding annotation databases were downloaded from Bioconductor. The visualization of GO enrichment was generated through *dotplot* function displaying the top 10 enriched biological process terms. As for calculating region-specific marker gene enrichment, we extracted gene expression from the marker gene and calculated the average gene expression within each predicted region. We then calculated the fold change of each marker gene in the expected region versus other regions as the enrichment score. Higher enrichment score indicates higher region detection accuracy. When calculating the significance of overlapping GO terms, we performed permutation test. We first unionized all GO terms, and, in each permutation, we sampled the GO terms according to each original numbers. Then, we calculated the number of overlapping GO terms and constructed a distribution. Then, our observed values were computed as a Z-score and p-value was considered as a two-tailed, either very small overlap or large number of overlaps. When comparing differentially expressed genes (DEGs) from two domains, we used *FindMarkers* function from *Seurat* package and set one of the domains as baseline. We then considered those ones with adjusted p-value less than 0.05 and an absolute value of average log2 fold change larger than 1 as significantly up-/down-regulated genes. Then, with these genes, we performed GO analysis with *clusterProfiler* R package as well as cell type enrichment using enrichR^52^. For gene expression similarity, we performed Spearman correlation based on the mean of normalized counts and removed those genes with average expression lower than 0.5 following Axolotl study^41^.

## Data availability

All datasets are publicly available, and the access numbers or the downloading websites are provided in the original publications as well. The Mouse Organogenesis Dataset^93^ measured by 10X Visium is available in the GEO dataset under accession code GSE214989. The Chicken Cardiogenesis Dataset^45^ measured by 10X Visium is available in the GEO dataset under accession code GSE149457. The Axolotl Telecephalon Neurogenesis Dataset^41^ measured by Stereo-seq is available at https://db.cngb.org/stomics/artista/download/. The Mouse Whole Brain Dataset^8^ measured by Stereo-seq is available at https://db.cngb.org/stomics/mosta/download/. More details on data preprocessing were provided in the Methods section.

## Code availability

The SpaDOT software code is publicly available at https://github.com/marvinquiet/SpaDOT. Detailed tutorials on installation and usage are provided at https://marvinquiet.github.io/SpaDOT/. SpaDOT is built as a Python package, and the recommended version is v3.9.18. There are several Python package dependencies, including torch (v2.0.1), torch_geometric (v2.6.1), anndata (v0.9.1), scanpy (v1.9.8), numpy (v1.22.4), pandas (v1.3.5), sklearn (v1.3.0) and rpy2 (v3.5.17). R environment (v4.3.3) and R package SPARK (https://github.com/xzhoulab/SPARK) are also recommended to be installed for spatially variable genes selection.

## Acknowledgements

This study was supported by the National Institutes of Health (NIH) Grant R01HG009124, R01GM144960, R01GM126553 and R01HG011883.

## Author contributions

X.Z and W.M conceived the idea and X.Z. provided funding support. W.M. and X.Z. designed the experiments with input from S.H., L.S. and J.L. W.M. developed the method, implemented the software, performed simulation and analyzed real data. W.M. and X.Z. wrote the manuscript with input from S.H., L.S. and J.L.

## Supplementary information

**Supplementary file 1:** Supplementary Note 1. Supplementary Tables S1-S2. Supplementary Figures S1-S24.

