## Supplementary File for "Optimal transport modeling uncovers spatial domain dynamics in spatiotemporal transcriptomics studies"

**This PDF file includes:**

Supplementary Note 1

Supplementary Tables S1-S2

Supplementary Figures S1-S24

References

### Supplementary Note 1

Traditional VAEs assume the latent variables $\boldsymbol{Z}$ to follow a multivariate normal distribution with an identity covariance matrix and the evidence lower bound (ELBO) is:

$$\mathrm{ELBO}_{\mathrm{VAE}} = \mathbb{E}_{q_\theta(\boldsymbol{Z}|\boldsymbol{Y})}[\log p_\psi(\widetilde{\boldsymbol{Y}}|\boldsymbol{Z})] - \beta \mathrm{D}_{\mathrm{KL}}(q_\theta(\boldsymbol{Z}|\boldsymbol{Y}) \parallel p(\boldsymbol{Z}))$$

where $p(\boldsymbol{Z}) \sim \mathcal{N}(\boldsymbol{Z}|0, \mathbf{I})$ implying orthogonality among the latent variables, $q_\theta$ denotes the encoder network parameterized by $\theta$, and $p_\psi$ denotes the decoder network parameterized by $\psi$. In $\mathrm{ELBO}_{\mathrm{VAE}}$, the first term denotes the reconstruction loss that measures how well the model reconstructs the original data from the latent space and the second term denotes the Kullback-Leibler (KL) divergence between the approximated posterior and a prior. We modify the traditional VAE model to incorporate spatial coordinates in two ways while sharing a common decoder. The reconstruction loss then involves two reconstruction terms: $\mathbb{E}_{q_{\theta_1}(\boldsymbol{Z_1}|\boldsymbol{Y},\boldsymbol{S})}[\log p_\psi(\widetilde{\boldsymbol{Y}}|\boldsymbol{Z_1})] + \mathbb{E}_{q_{\theta_2}(\boldsymbol{Z_2}|\boldsymbol{Y})}[\log p_\psi(\widetilde{\boldsymbol{Y}}|\boldsymbol{Z_2})]$, where $\boldsymbol{Z_1}$ has conditional prior given spatial locations $\boldsymbol{S}$. Since we assume $\boldsymbol{Z_1}$ and $\boldsymbol{Z_2}$ to be independent, we concatenate their latent spaces and redefine the reconstruction loss as $\mathbb{E}_{q(\boldsymbol{Z_1} \oplus \boldsymbol{Z_2}|\boldsymbol{Y},\boldsymbol{S})}[\log p_\psi(\widetilde{\boldsymbol{Y}}|\boldsymbol{Z_1} \oplus \boldsymbol{Z_2})]$ and $\oplus$ denotes concatenation. Details of the encoders and KL terms are separately stated in the following paragraphs.

**SVGPVAE details.** The goal of Gaussian Process Variational Autoencoder (GPVAE) is to train a model that can generate spatial gene expression $\boldsymbol{Y}$ conditioned on spatial coordinates $\boldsymbol{S}$ while simultaneously inferring an interpretable and disentangled latent space. To achieve this, the prior on the latent space $\boldsymbol{Z_1}$ conditioned on $\boldsymbol{S}$ is defined as $p(\boldsymbol{Z_1}|\boldsymbol{S}) = \prod_{d_1} \mathcal{N}(\boldsymbol{z_1}^{d_1}{}_{1:n} \,|\, 0, \mathbf{K}(\boldsymbol{S}, \boldsymbol{S}))$ where the covariance function $\mathbf{K}(\boldsymbol{S}, \boldsymbol{S})$ denotes the correlation of different channels in the latent space and captures spatial dependencies. Then, based on the latent space $\boldsymbol{Z_1}$, the generative process of $\boldsymbol{Y}$ can be denoted as $p(\boldsymbol{Y}|\boldsymbol{Z_1}) = \prod_n \mathcal{N}\left(\boldsymbol{y}_n \middle| \mu_\psi(\boldsymbol{z}_{1_n}), \sigma_\psi^2(\boldsymbol{z}_{1_n})\mathbf{I}\right)$ where $\mu_\psi(\cdot)$ and $\sigma_\psi(\cdot)$ denote the set of parameters from the decoder. The joint distribution becomes $p(\boldsymbol{Y}, \boldsymbol{Z_1}|\boldsymbol{S}) = p(\boldsymbol{Y}|\boldsymbol{Z_1})p(\boldsymbol{Z_1}|\boldsymbol{S})$ and the derivation of $\boldsymbol{Z_1}$ becomes $p(\boldsymbol{Z_1}|\boldsymbol{Y}, \boldsymbol{S}) = \frac{p(\boldsymbol{Y}, \boldsymbol{Z_1}|\boldsymbol{S})}{p(\boldsymbol{Y}|\boldsymbol{S})} = \frac{p(\boldsymbol{Y}|\boldsymbol{Z_1})p(\boldsymbol{Z_1}|\boldsymbol{S})}{p(\boldsymbol{Y}|\boldsymbol{S})}$. Considering the intractable property of $\boldsymbol{Z_1}$ and intractable likelihood $p(\boldsymbol{Y}|\boldsymbol{Z_1})$, Pearce (2020)[1] proposes to use an inference network (encoder) parameterized by $\theta_1$: $\tilde{q}_{\theta_1}(\boldsymbol{Z_1}|\boldsymbol{Y}) = \prod_n \mathcal{N}\left(\boldsymbol{z}_{\mathbf{1}_n} \middle| \mu_{\theta_1}(\boldsymbol{y}_n), \text{diag}\left(\sigma_{\theta_1}^2(\boldsymbol{y}_n)\right)\right)$ to replace $p(\boldsymbol{Y}|\boldsymbol{Z_1})$ in the posterior which would result in $q(\boldsymbol{Z}|\boldsymbol{Y}, \boldsymbol{S}) \propto \tilde{q}_{\theta_1}(\boldsymbol{Z_1}|\boldsymbol{Y})p(\boldsymbol{Z_1}|\boldsymbol{S}) = \prod_{d_1}\prod_n \mathcal{N}\left(\boldsymbol{z}_{\mathbf{1}_n}^{d_1} \middle| \mu_{\theta_1}^{d_1}(\boldsymbol{y}_n), \text{diag}\left(\sigma_{\theta_1}^{d_1}(\boldsymbol{y}_n)^2\right)\right)\mathcal{N}(\boldsymbol{z}_{\mathbf{1}}^{d_1}{}_{1:n}|\, 0, \mathbf{K}(\boldsymbol{S}, \boldsymbol{S}))$. Due to the symmetrical property in the probability density function of Gaussian distribution, $\mathcal{N}(z|\mu, \sigma) = \mathcal{N}(\mu|z, \sigma)$, the approximate posterior for each channel becomes the exact GP posterior in traditional GP regression. In this context, the input is the spatial coordinates $\boldsymbol{S}$ and output corresponds to encoder output $\mu_{\theta_1}(\boldsymbol{Y})$ with heteroscedastic noise $\sigma_{\theta_1}(\boldsymbol{Y})^2$.

However, when dealing with a large amount of data, calculating the exact GP posterior may encounter scalability issues. To ensure the model's scalability, a sparse variational GP[2] is employed to approximate the full GP using by strategically choosing a set of inducing points $\boldsymbol{U}$ from $\boldsymbol{S}$ with corresponding inducing variables $u = f(\boldsymbol{U})$, where $p(u) \sim \mathcal{N}\left(0, \mathbf{K}(\boldsymbol{U}, \boldsymbol{U})\right)$. Then, the original posterior $p\left(f \middle| \boldsymbol{S}, \boldsymbol{z_1}_{1:n}^{d_1}\right)$ is approximated by $\int p(f|u)p\left(u \middle| \boldsymbol{S}, \boldsymbol{z_1}_{1:n}^{d_1}\right) \mathrm{d}u$ and the posterior distribution of $u$ is given by $p\left(u \middle| \boldsymbol{S}, \boldsymbol{z_1}_{1:n}^{d_1}\right) \propto p\left(\boldsymbol{z_1}_{1:n}^{d_1} \middle| u\right)p(u)$. One of the most popular approaches is using variational inference to approximate the true posterior with a simpler variational distribution $q(u) = \mathcal{N}(\boldsymbol{\mu}_u, \boldsymbol{A}_u)$ and we therefore have $\mathbf{L}_{\text{SVGPVAE-KL}} = \mathbb{E}_{q(f)}\left[\log p\left(\boldsymbol{z}_{\mathbf{1}}{}_{1:n}^{d_1} | f\right)\right] - \mathrm{D}_{\mathrm{KL}}\left(q(u) \parallel p(u)\right)$.

**GATVAE details**. The goal of GATVAE is to map the graph $G$ constructed on spatial coordinates $\boldsymbol{S}$ into a latent space $\boldsymbol{Z}_{\mathbf{2}}$[3,4]. Here, we construct graph $G$ based on K-Nearest Neighbor and generate an adjacency matrix $A$ where $A_{ij} = 1$ indicates node $i$ and node $j$ are neighbors and $A_{ij} = 0$ otherwise. We again approximate the true posterior with $q_{\theta_2}(\boldsymbol{Z}_{\mathbf{2}} | \boldsymbol{Y}, \boldsymbol{A})$ where $\boldsymbol{Z}_{\mathbf{2}} = \left[\boldsymbol{z}_{\mathbf{2}_1}, \boldsymbol{z}_{\mathbf{2}_2}, \dots, \boldsymbol{z}_{\mathbf{2}_n}\right]$ and $\boldsymbol{z}_{\mathbf{2}_i}$ is a vector of size $d_2$ , denoting the latent space of node $i$. We consider the node features as initial node representations $\boldsymbol{z}_{\mathbf{2}_i} = \boldsymbol{y}_i$ and the $h^{th}$ encoder layer generates the representation of node $i$ in layer $h$:

$$\mathbf{z}_{\mathbf{2}_i}^{(h)} = \sum_{j \in \{\mathrm{N}(i) \cup i\}} \alpha_{ij}^{(h)} \, \sigma(\mathbf{W}^{(h)} \mathbf{z}_{\mathbf{2}_j}^{(h-1)})$$

where $\mathrm{N}(i)$ denotes the neighborhood of node $i$, $\alpha_{ij}^{(h)}$ is the attention coefficients from neighboring node $j$ to node $i$ in the $h^{th}$ encoder layer, $\mathbf{W}^{(h)}$ is a trainable parameter of $h^{th}$ encoder layer and $\sigma(\cdot)$ denotes the activation function. In each layer, the attention coefficients are normalized from the attention relevance $e_{ij}$ by using the softmax function: $\alpha_{ij}^{(h)} = \frac{\exp\left(e_{ij}^{(h)}\right)}{\sum_{k \in \{\mathrm{N}(i) \cup i\}} \exp\left(e_{ik}^{(h)}\right)}$ . The relevance $e_{ij}^{(h)}$ is computed by $\text{LeakyRELU}(\mathbf{v}^{(h)^T} (\mathbf{W}^{(h)} \mathbf{z}_{\mathbf{2}_i}^{(h-1)} \oplus \mathbf{W}^{(h)} \mathbf{z}_{\mathbf{2}_j}^{(h-1)}))$ and $\oplus$ denotes the concatenation of two latent space and $\mathbf{v}^{(h)}$ is another trainable parameter of $h^{th}$ encoder layer.

**Implementation details**. For the decoder, we utilize a multi-layer perceptron (MLP) architecture consisting of two hidden layers, containing 64 nodes and 256 nodes respectively. Each layer is followed by layer normalization and Exponential Linear Unit (ELU) as activation function. The encoder for SVGPVAE also adopts an MLP architecture with two hidden layers containing 256 and 64 nodes, respectively. Each layer is followed by batch normalization and the ELU activation function. For SVGPVAE encoder, we utilize 1,200 inducing points in total and proportionally assign the number of inducing points for each time point based on their number of spots / cells. We fix the inducing points once we sample them at the beginning phase of the training and use a fixed bandwidth for kernel function. For GATVAE encoder, we use three layers each with hidden node size of 512 and number of attention heads as 4. In the first two layers, the multi-head attentions are concatenated while in the final layer, they are averaged. We use ELU activation function for the first two layers.

### Supplementary Tables

Supplementary Table S1. Summary of model parameters used in SpaDOT. This table includes all fixed and tunable parameters. Default values are listed unless otherwise specified.

| Parameter Name | Symbol | Description | Value |
|---|---|---|---|
| Latent dimension for SVGP encoder | $\boldsymbol{Z_1}$ | Dimension of latent space capturing spatial gene expression patterns derived by SVGP encoder | 10 |
| Latent dimension for GAT encoder | $\boldsymbol{Z_2}$ | Dimension of latent space capturing spatial gene expression patterns derived by GAT encoder | 10 |
| Hidden layer structure of encoder | - | Number of neurons in hidden layers of encoder | [256, 64] |
| Hidden layer structure of decoder | - | Number of neurons in hidden layers of decoder | [64, 256] |
| Learning rate | - | Learning rate for AdamW | 3e-4 |
| Batch size | - | Number of spatial spots / cells per training batch | 512 |
| Number of epochs | - | Total training epochs | 100 |
| Number of epochs adding OT loss | - | Number of training epochs when introducing OT constraints | 50 |
| Reconstruction loss of VAE | - | Weight for reconstruction loss | 0.1 |
| KL divergence weight of SVGP | $\beta_1$ | Weight for KL loss $\mathbf{L}_{\mathrm{SVGPVAE-KL}}$ | 1 |
| KL divergence weight of GAT | $\beta_2$ | Weight for KL loss $\mathbf{L}_{\mathrm{GATVAE-KL}}$ | 1e-4 |
| Alignment weight | $\omega_1$ | Weight for alignment loss $\mathbf{L}_{\mathrm{Alignment}}$ | 0.1 |
| K-Means weight | $\omega_2$ | Weight for K-Means loss $\mathbf{L}_{\mathrm{KMeans}}$ | 0.1 |
| OT weight | $\omega_3$ | Weight for OT loss $\mathbf{L}_{\mathrm{OT}}$ | 1 |
| No. clusters during training | $k$ | Number of clusters during training to impose clustering constraints | 10 |
| No. inducing points | $u$ | Number of inducing points sampled for samples collected from all time points | 1,200 |
| Kernel type | - | The choice of GP kernel | RBF |
| Lengthscale of RBF kernel | $l$ | The length scale of the SVGP RBF kernel | 0.1 |

* SVGP: stochastic variational gaussian process; GAT: graph attention transformer; VAE: variational autoencoder; KL: Kullback–Leibler; OT: optimal transport; RBF: radial basis function.

* For parameters in optimal transport (OT), we use default parameters in Waddington-OT [5] but setting $\lambda_1 = 0.1, \lambda_2 = 5$.

Supplementary Table S2. Comparison of strategies between SpaDOT and other competing methods.

| **Methods** | **Designed for multi-slices?** | **Relaxation on number of clusters across multi-slices?** | **Default clustering methods** |
|---|---|---|---|
| spaDOT | ✓ | ✓ | K-Means |
| SpatialPCA | ✗ | ✓ | Walktrap |
| GraphST | ✓ | ✗ | Mclust |
| SEDR | ✓ | ✗ | Mclust |
| spaGCN | ✗ | ✓ | Iterative clustering with Louvain |
| spaVAE | ✓ | ✓ | K-Means / Louvain |
| STAligner | ✓ | ✗ | Mclust |
| PRECAST | ✓ | ✗ | Potts model |
| spatiAlign | ✓ | ✗ | Hierarchical clustering |
| SPIRAL | ✓ | ✗ | Mclust |
| STADIA | ✓ | ✗ | Gaussian Mixture model |

### Supplementary Figures

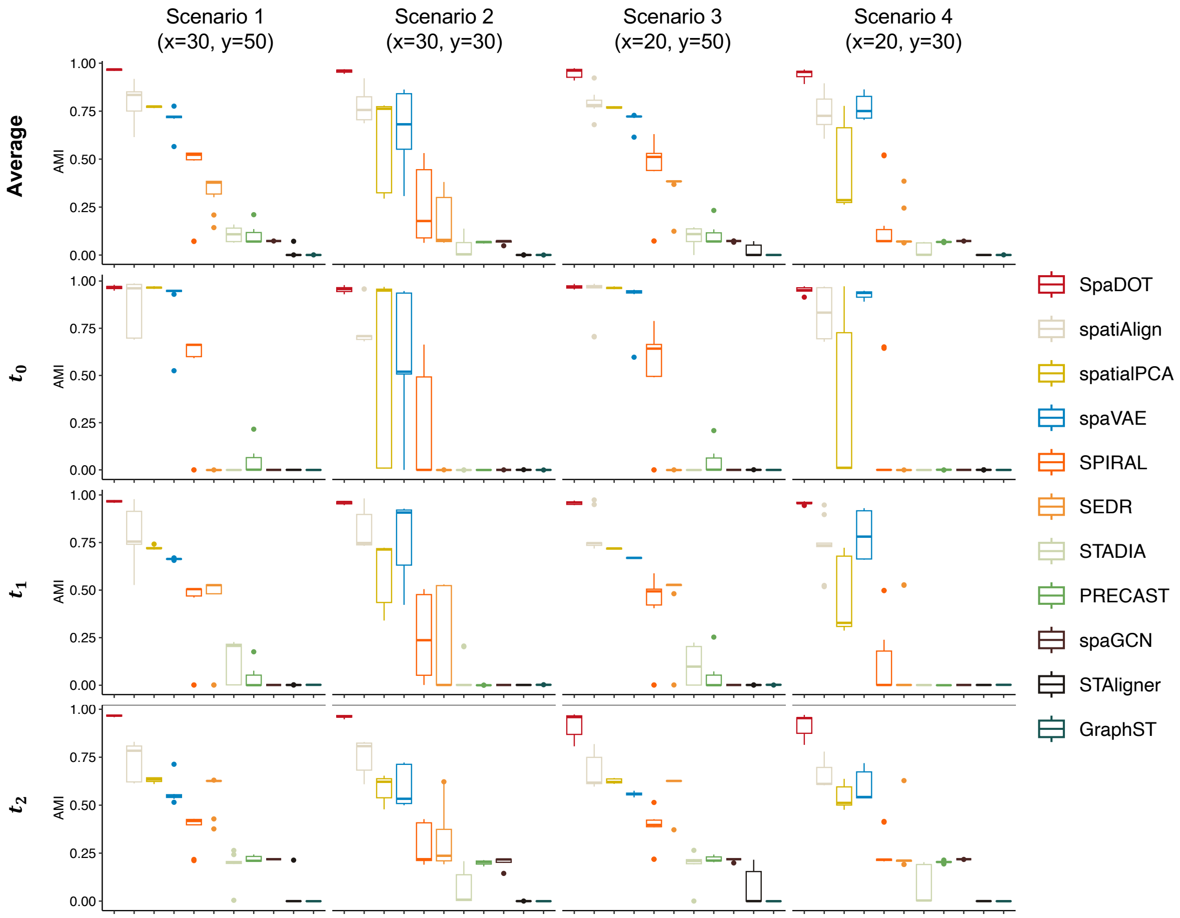


**Supplementary Figure S1.** We evaluate domain detection accuracy using adjusted Mutual Information (AMI) and report AMI-avg across three time points, as well as per time point. Each box in the boxplots represents ten replicates per method. In the boxplot, the center line, box limits and whiskers denote the median, upper and lower quartiles, and 1.5× interquartile range.

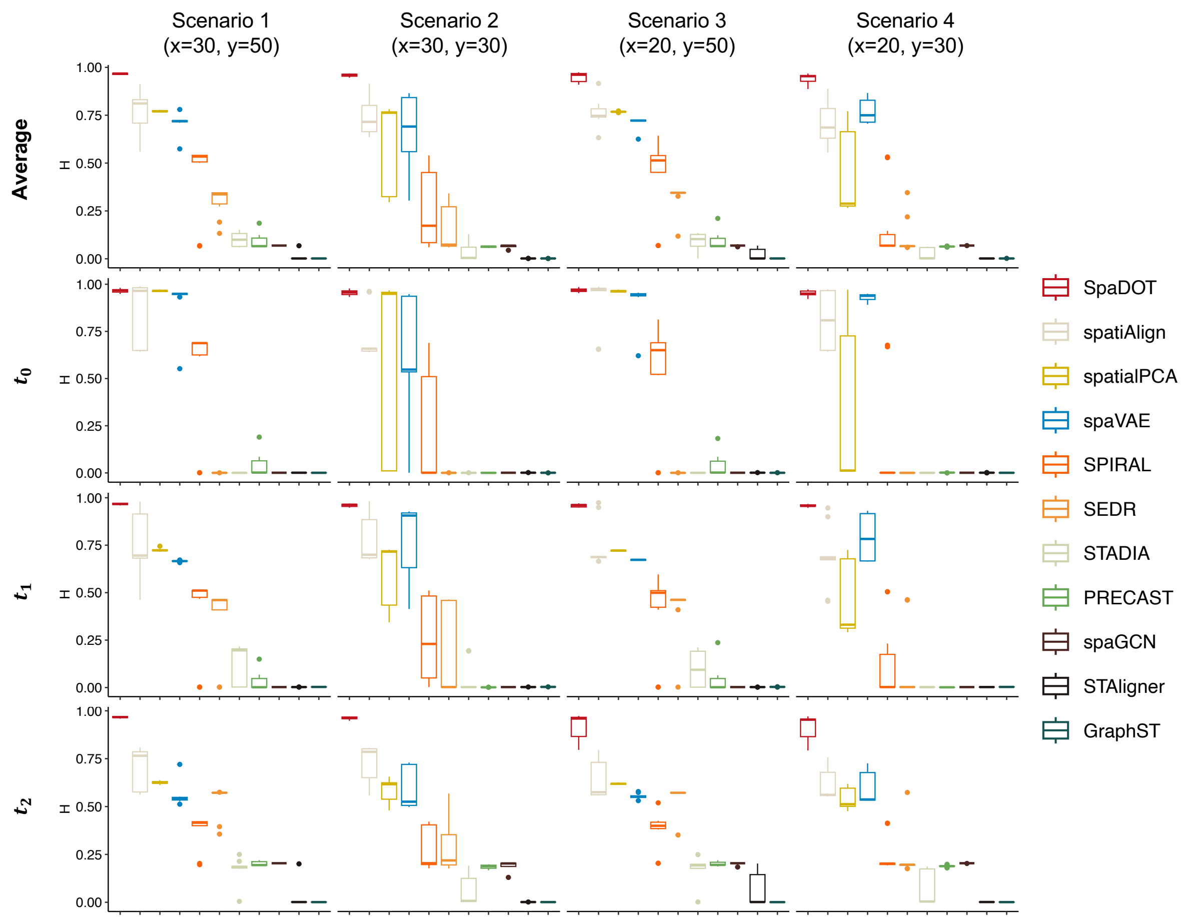


**Supplementary Figure S2.** We evaluate domain detection accuracy using Homogeneity (H) and report H-avg across three time points, as well as per time point. Each box in the boxplots represents ten replicates per method. In the boxplot, the center line, box limits and whiskers denote the median, upper and lower quartiles, and 1.5× interquartile range.

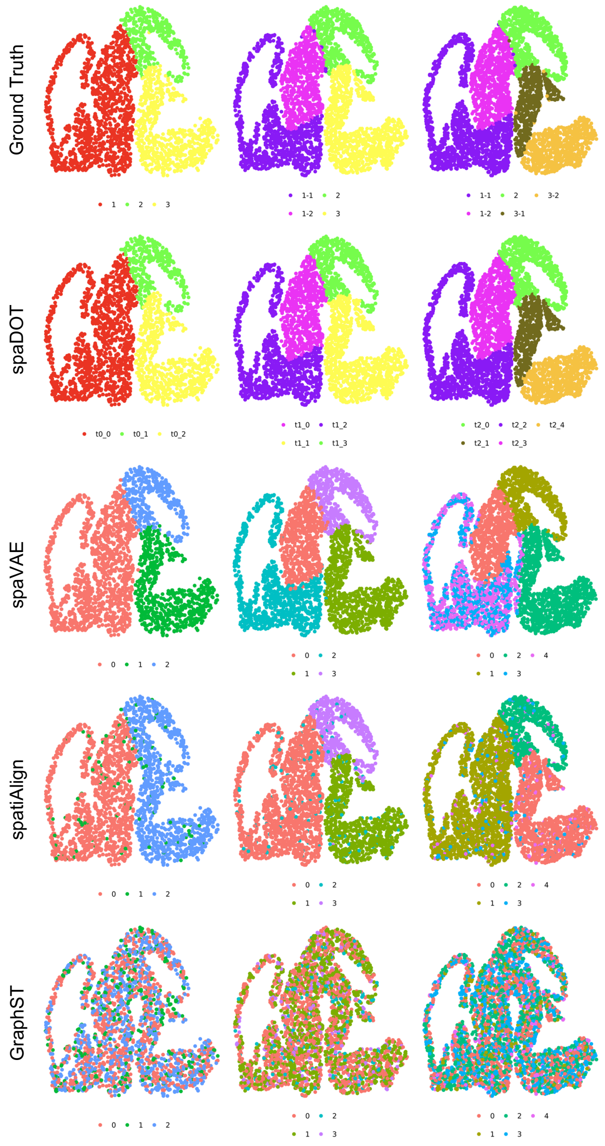


**Supplementary Figure S3.** Spatial domain layouts of ground truth and predicted domains by SpaDOT, spaVAE, spatiAlign, and GraphST. The colors used in SpaDOT predictions are curated for visual clarity.

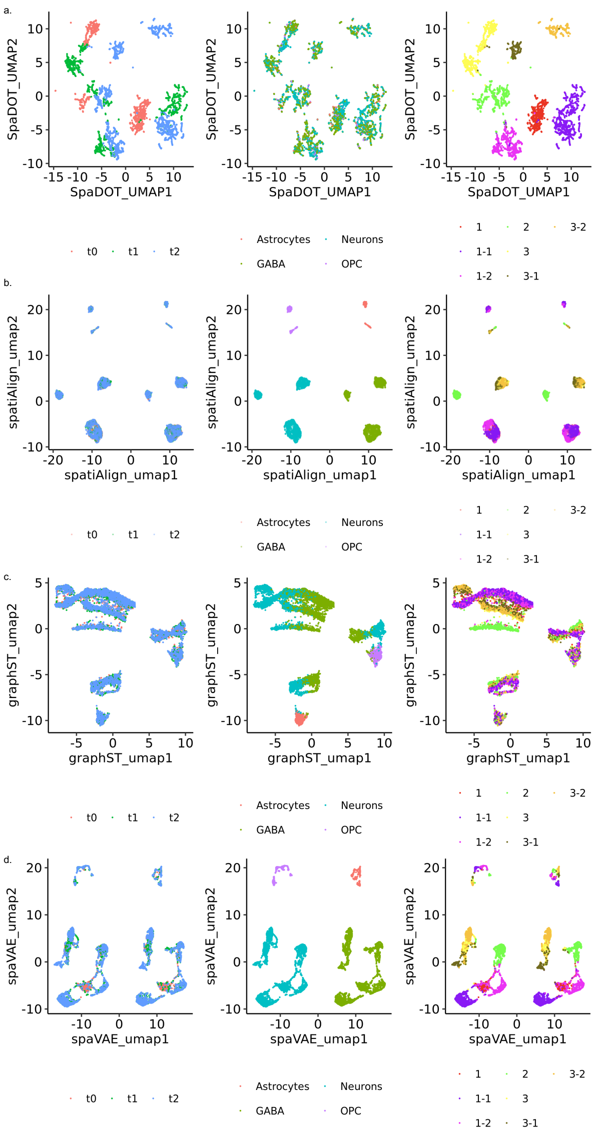


**Supplementary Figure S4.** UMAP visualization of latent representations learned by a) SpaDOT, b) spatiALign, c) GraphST, and d) spaVAE. For each method, the left panels are colored by time point, the middle panels by cell type, and the right panels by spatial domains.

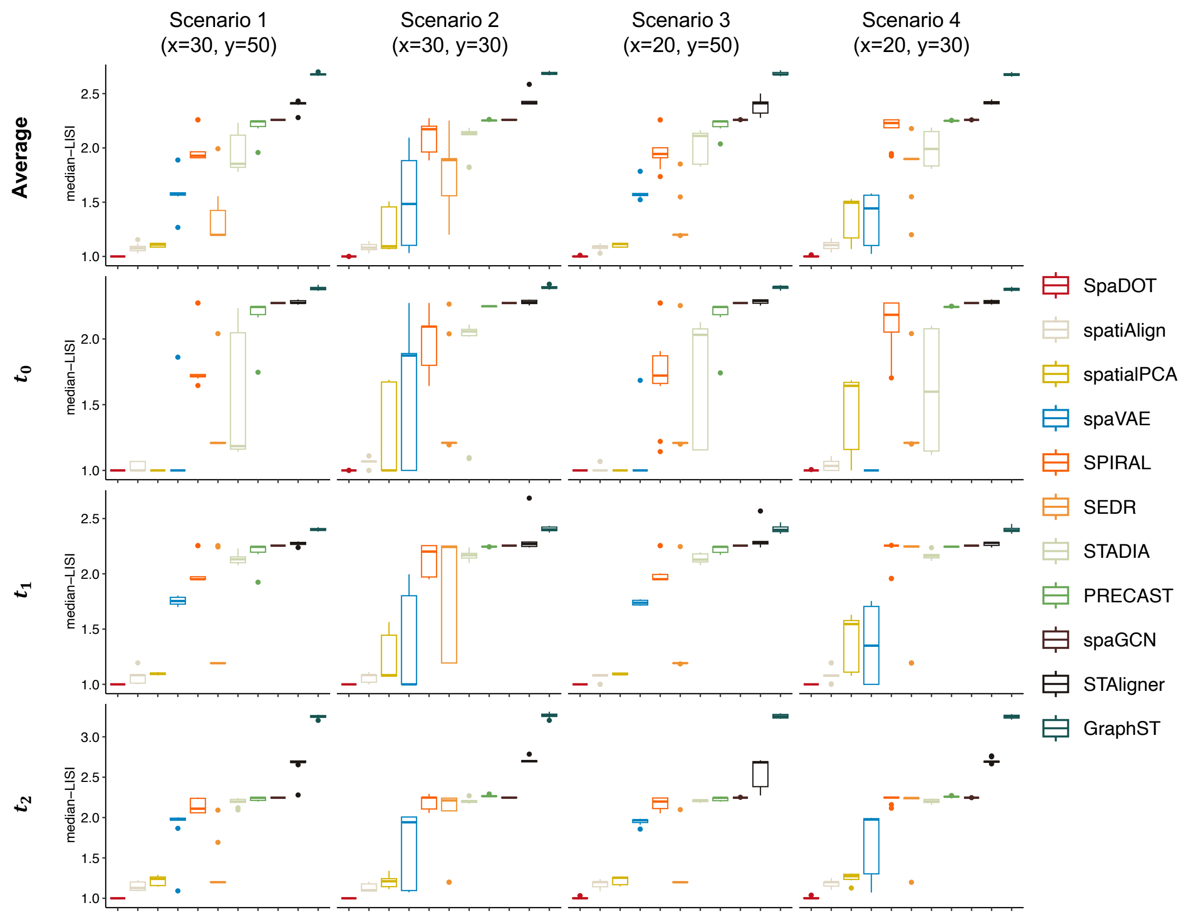


**Supplementary Figure S5.** We evaluate spatial domain continuity using median local inverse Simpson index (LISI) and report medianLISI-avg across three time points, as well as per time point. Each box in the boxplots represents ten replicates per method. In the boxplot, the center line, box limits and whiskers denote the median, upper and lower quartiles, and 1.5× interquartile range.

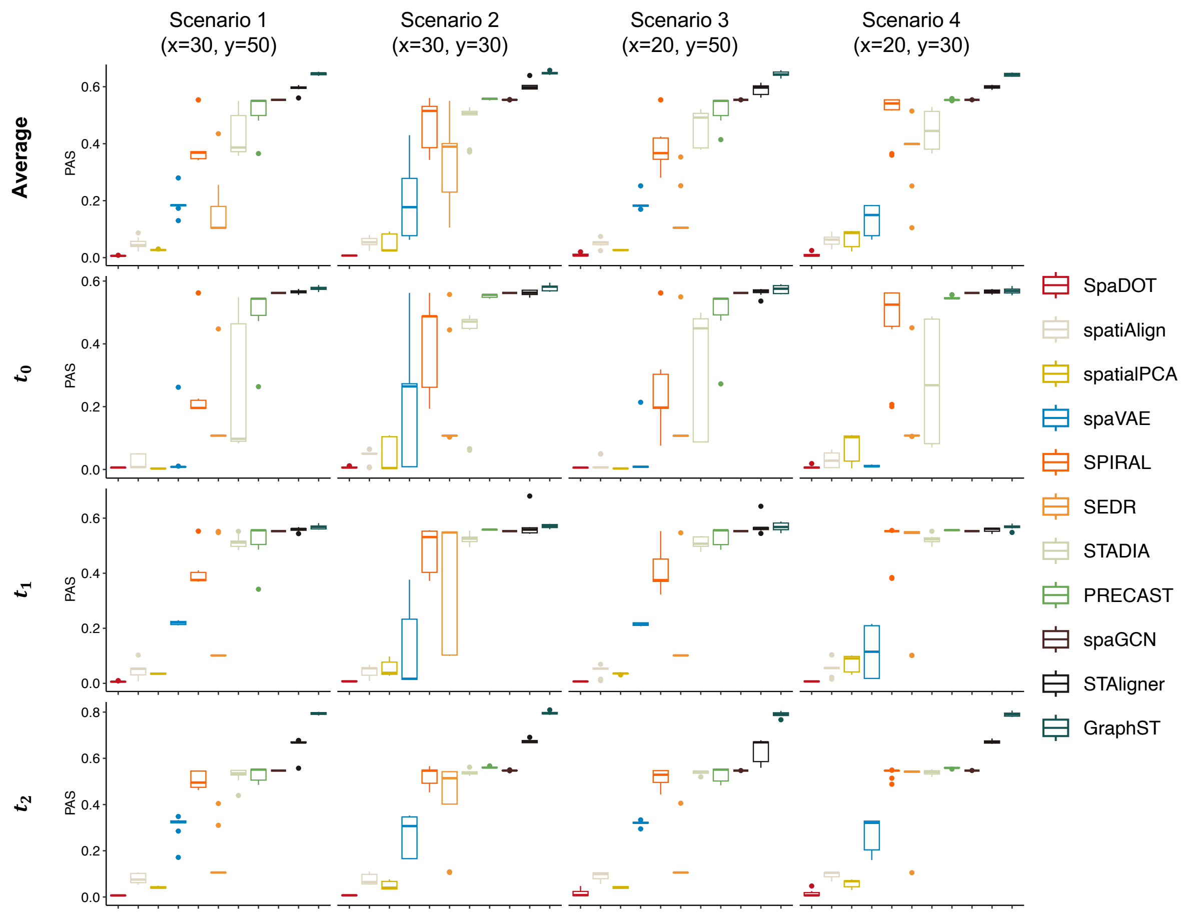


**Supplementary Figure S6.** We evaluate spatial domain continuity using percentage of abnormal spots (PAS) and report PAS-avg across three time points, as well as per time point. Each box in the boxplots represents ten replicates per method. In the boxplot, the center line, box limits and whiskers denote the median, upper and lower quartiles, and 1.5× interquartile range.

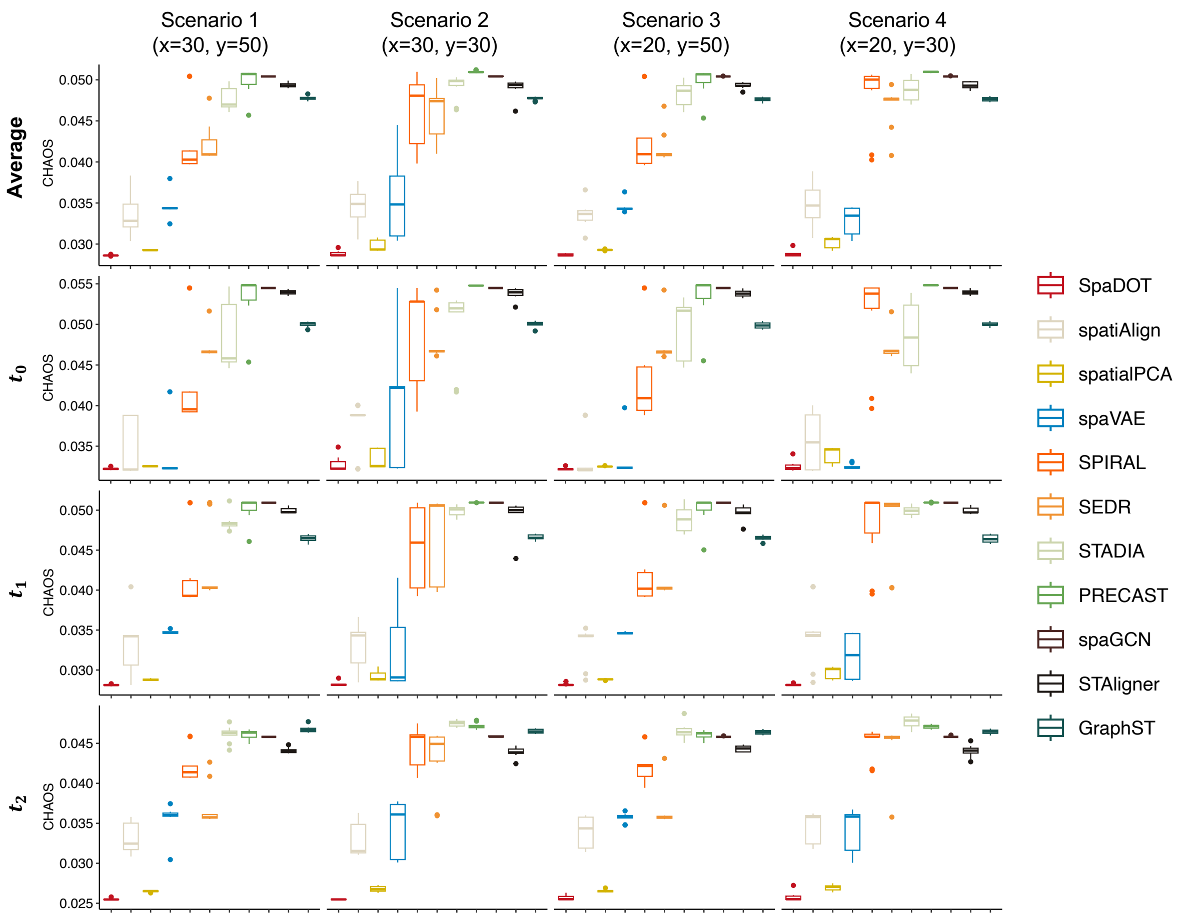


**Supplementary Figure S7.** We evaluate spatial domain continuity using spatial CHAOS score (CHAOS) and report CHAOS-avg across three time points, as well as per time point. Each box in the boxplots represents ten replicates per method. In the boxplot, the center line, box limits and whiskers denote the median, upper and lower quartiles, and 1.5× interquartile range.

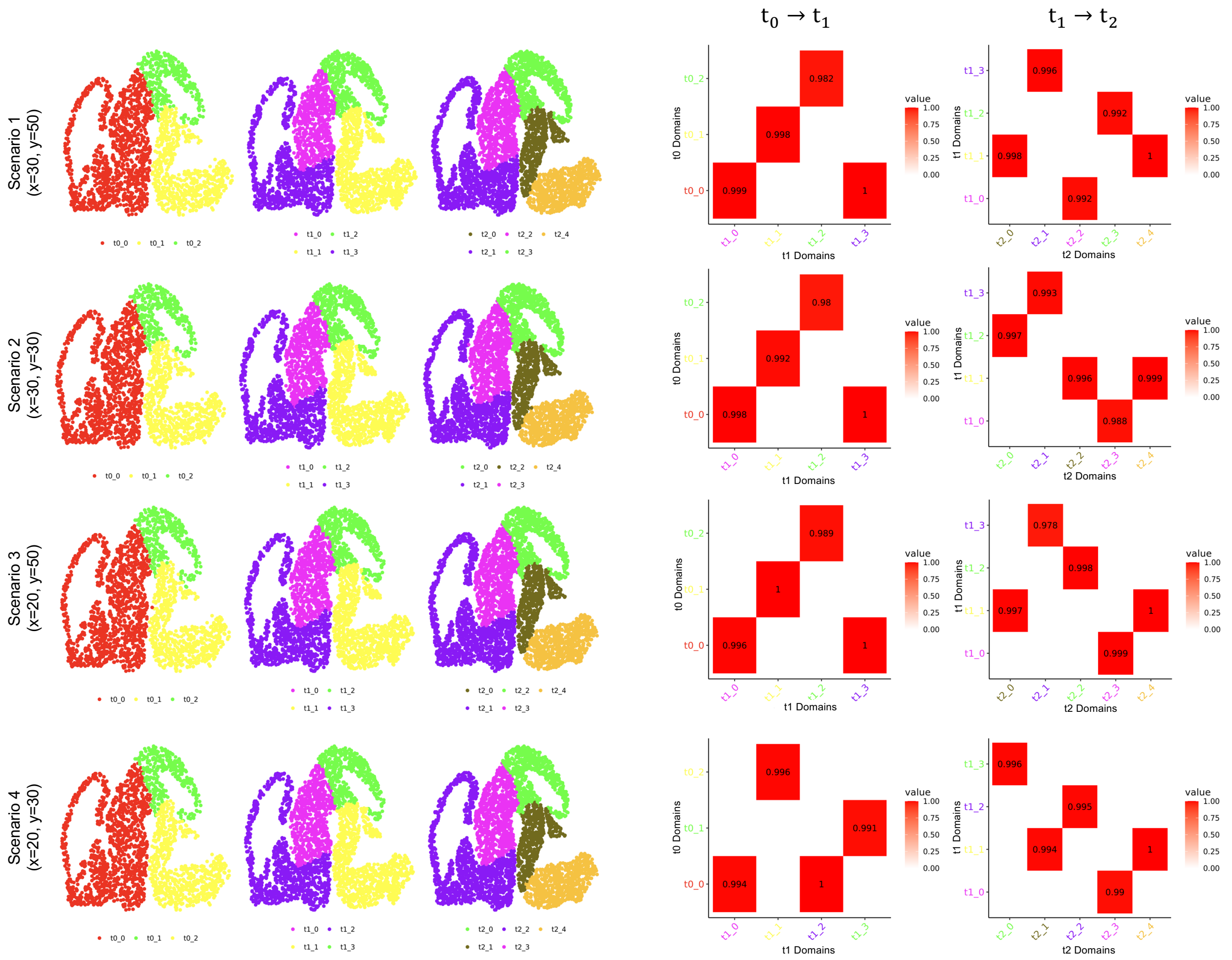


**Supplementary Figure S8.** Illustration of domain transition dynamics using one simulation replicate (seed: 1993) selected from ten replicates. From top to bottom are the four simulation scenarios. For each scenario, the left panels display the spatial layout of predicted domains across time points, while the right panels show the corresponding domain transition probabilities visualized as heatmaps.

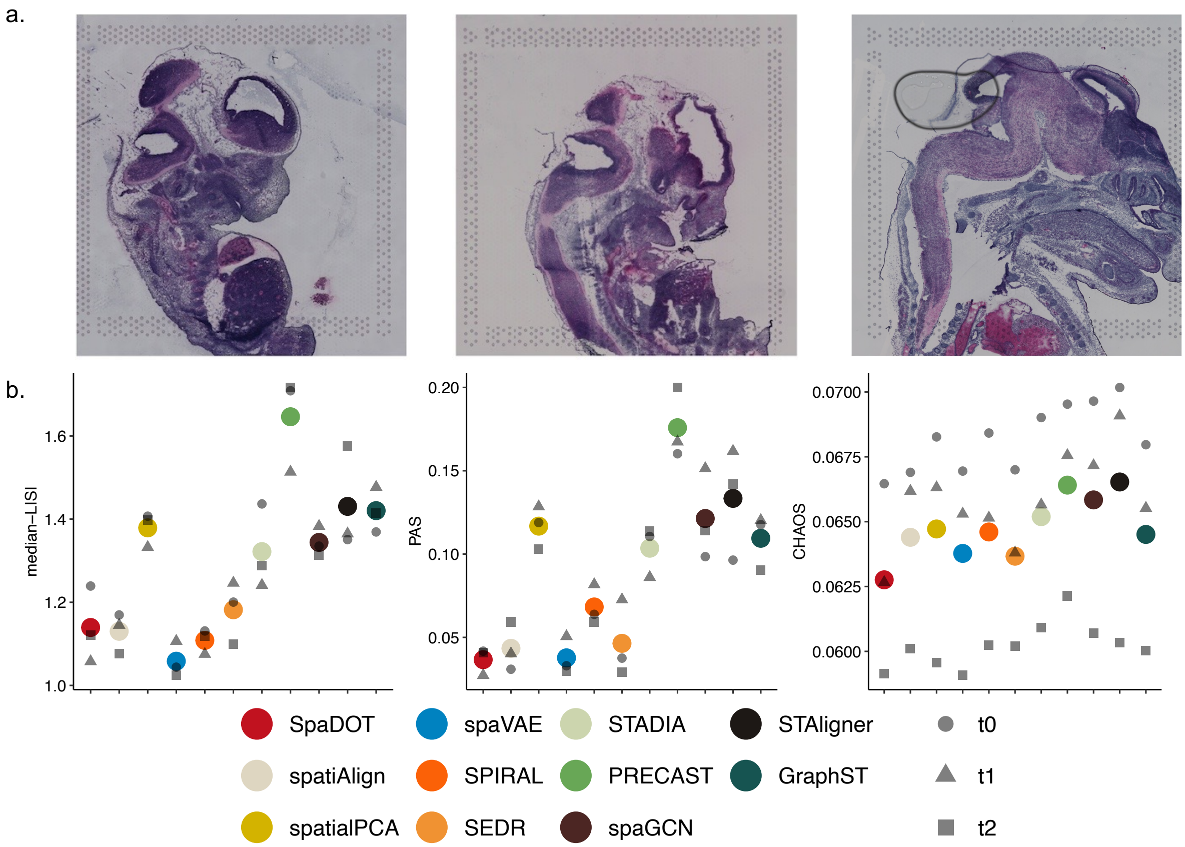


**Supplementary Figure S9.** a) H&E-stained histology images of mouse embryos at stages E12.5 (left), E13.5 (middle), and E15.5 (right). b) Spatial continuity evaluation using three metrics: median-LISI, PAS, and CHAOS. Colored dots indicate the average performance across all time points, while grey dots with different shapes represent performance at individual time points.

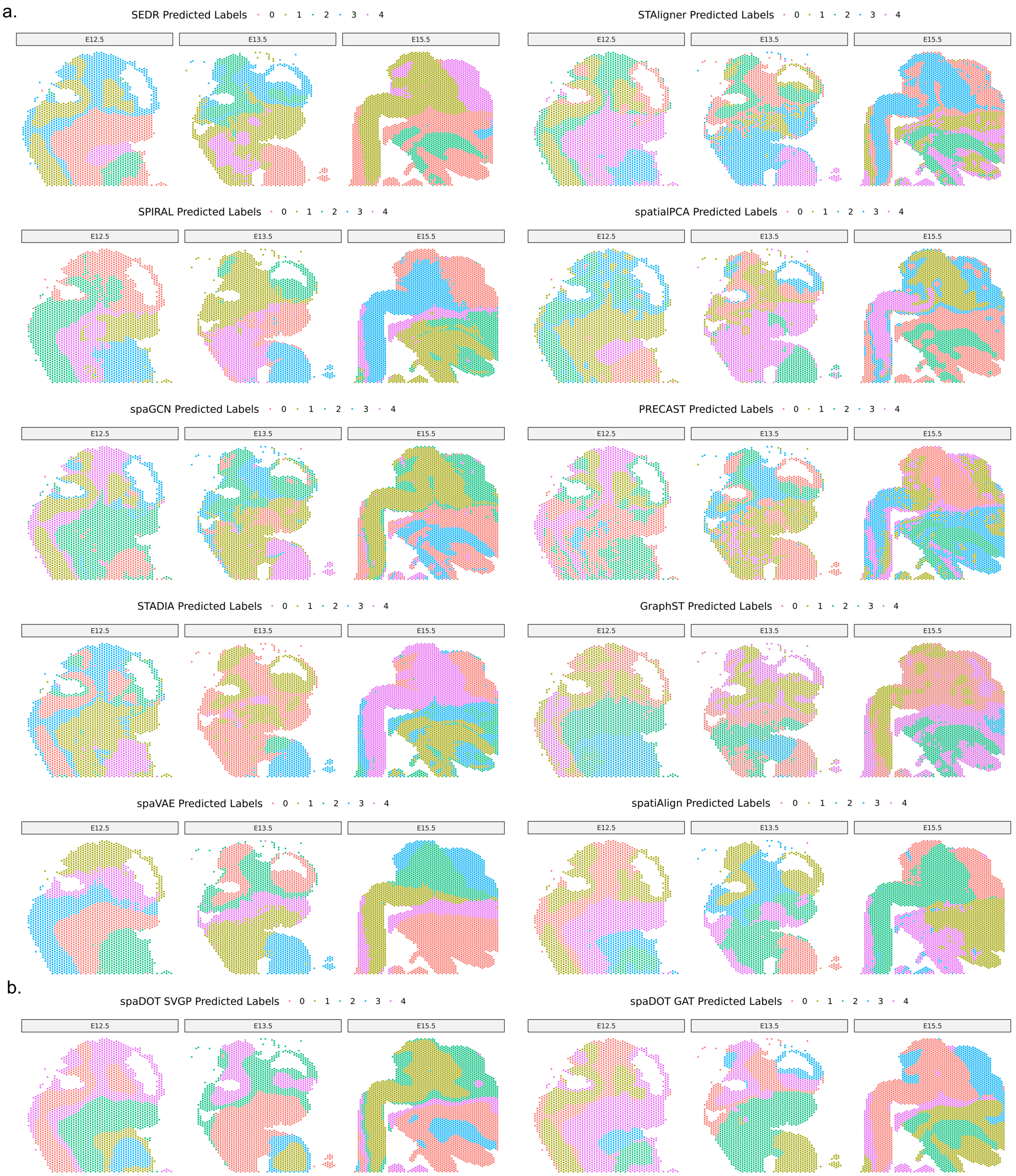


**Supplementary Figure S10.** a) Spatial domain layouts predicted by ten competing methods for comparison in mouse embryos. b) Spatial layouts generated using latent representations from individual components of SpaDOT: the left panel shows results from the stochastic variational Gaussian process (SVGP) encoder, and the right panel shows results from the graph attention transformer (GAT) encoder.

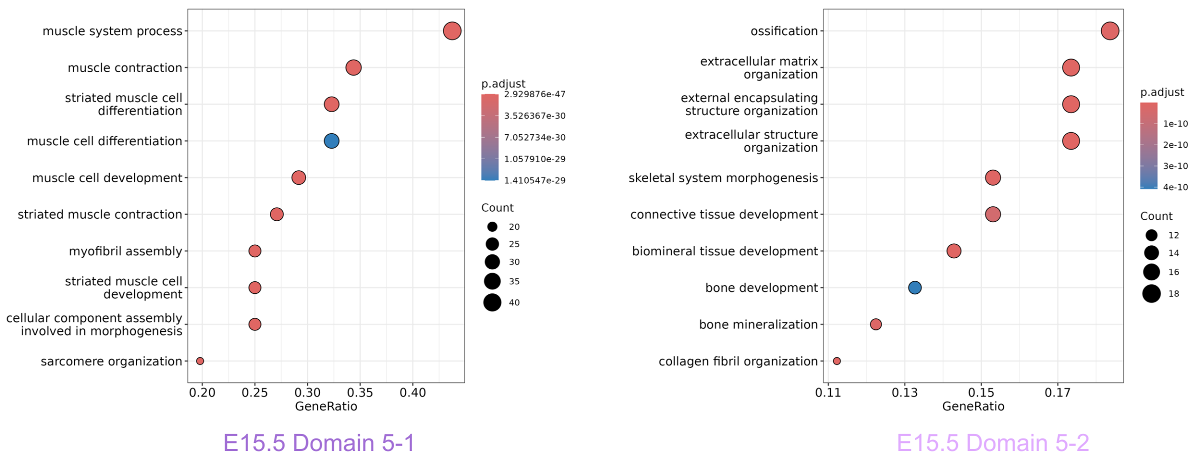


**Supplementary Figure S11.** Gene ontology (GO) enrichment analysis of domain-specific marker genes for domain 5-1 (left) and domain 5-2 (right) at embryonic stage E15.5.

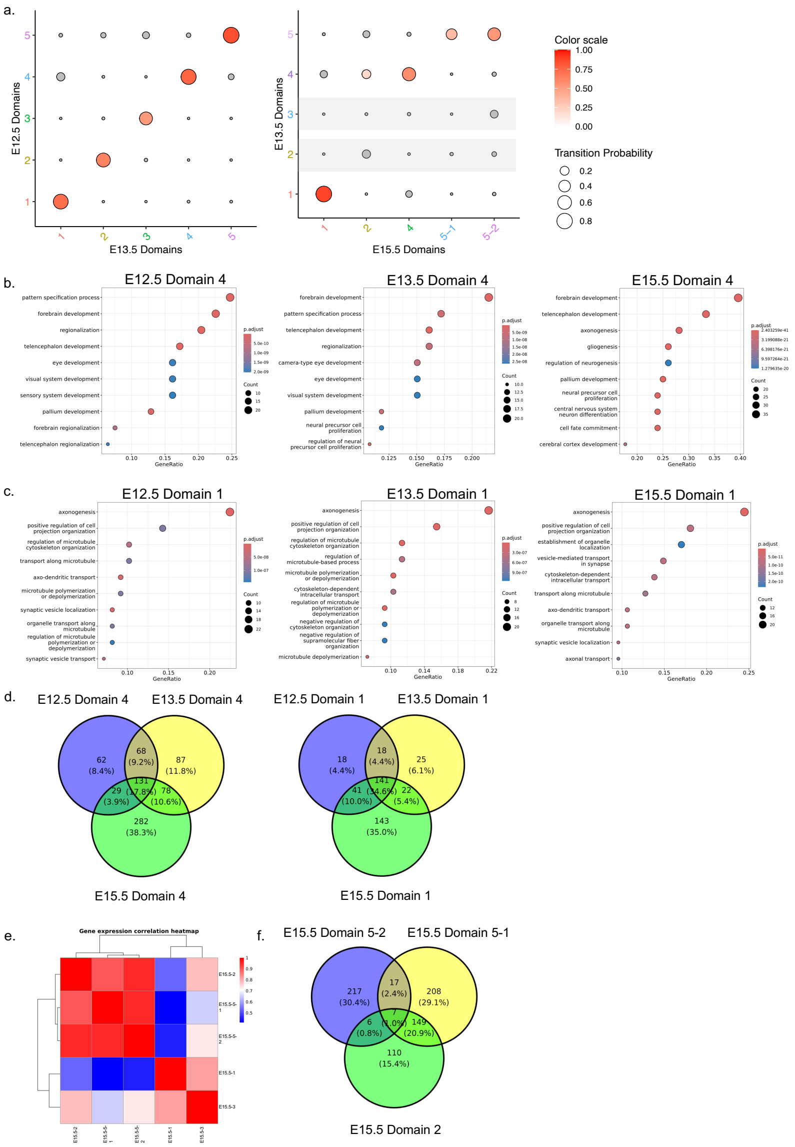


**Supplementary Figure S12.** a) Dot plot showing domain transition probabilities across time points. Shaded areas indicate no significant transitions with probability > 0.2. b) Gene ontology (GO) enrichment for domain 4 across all time points. c) Gene ontology (GO) enrichment for domain 1 across all time points. d) Venn diagrams showing the number of overlapping GO terms among domain 4 across all time points (left) and domain 1 across all time points (right). e) Heatmap shows Spearman correlation of averaged gene expressions among domains predicted at E15.5. f) GO term overlap between domains 2, 5-1, and 5-2 at E15.5.

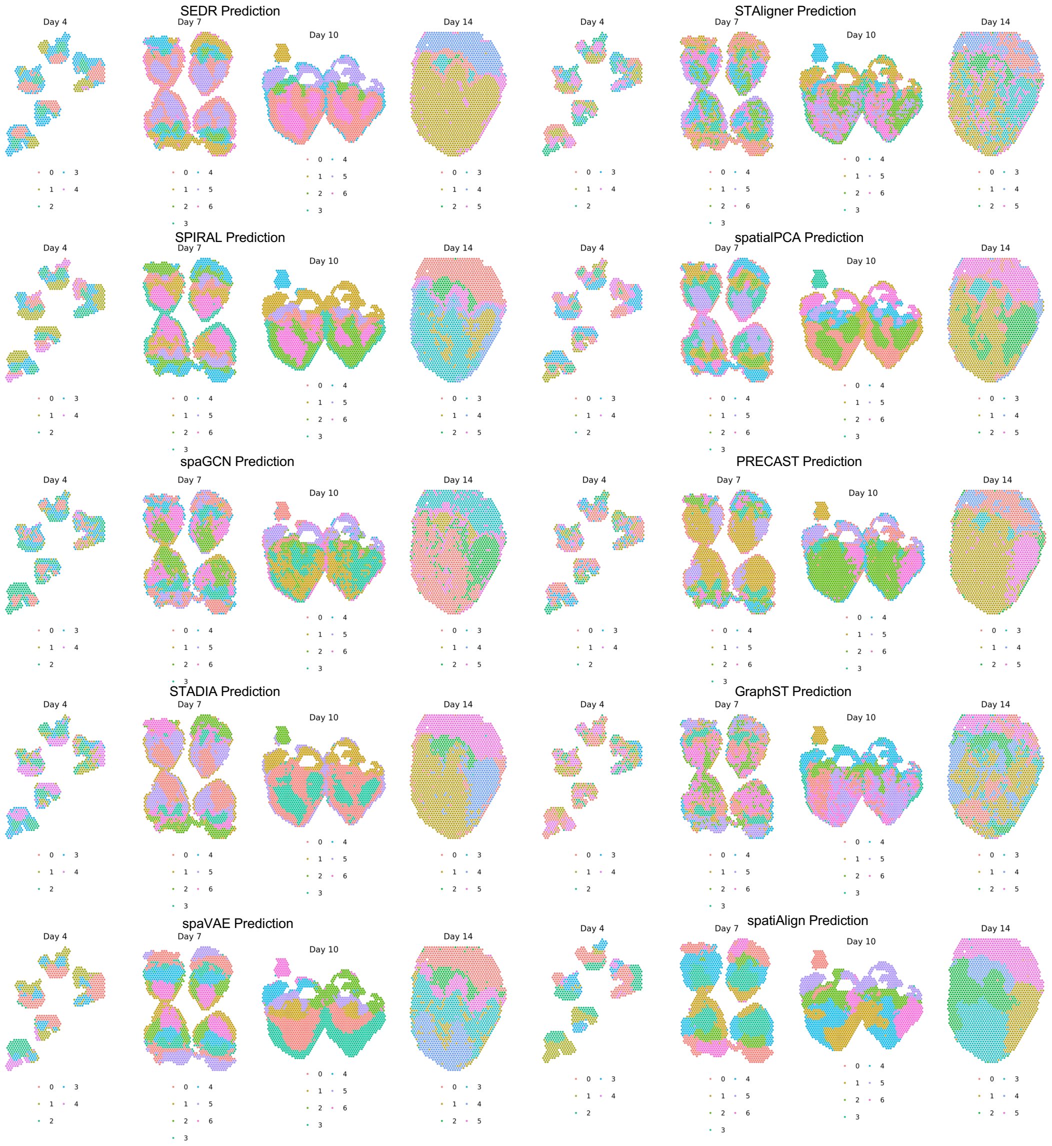


**Supplementary Figure S13.** Spatial domain layouts predicted by ten competing methods for comparison in developing chicken heart.

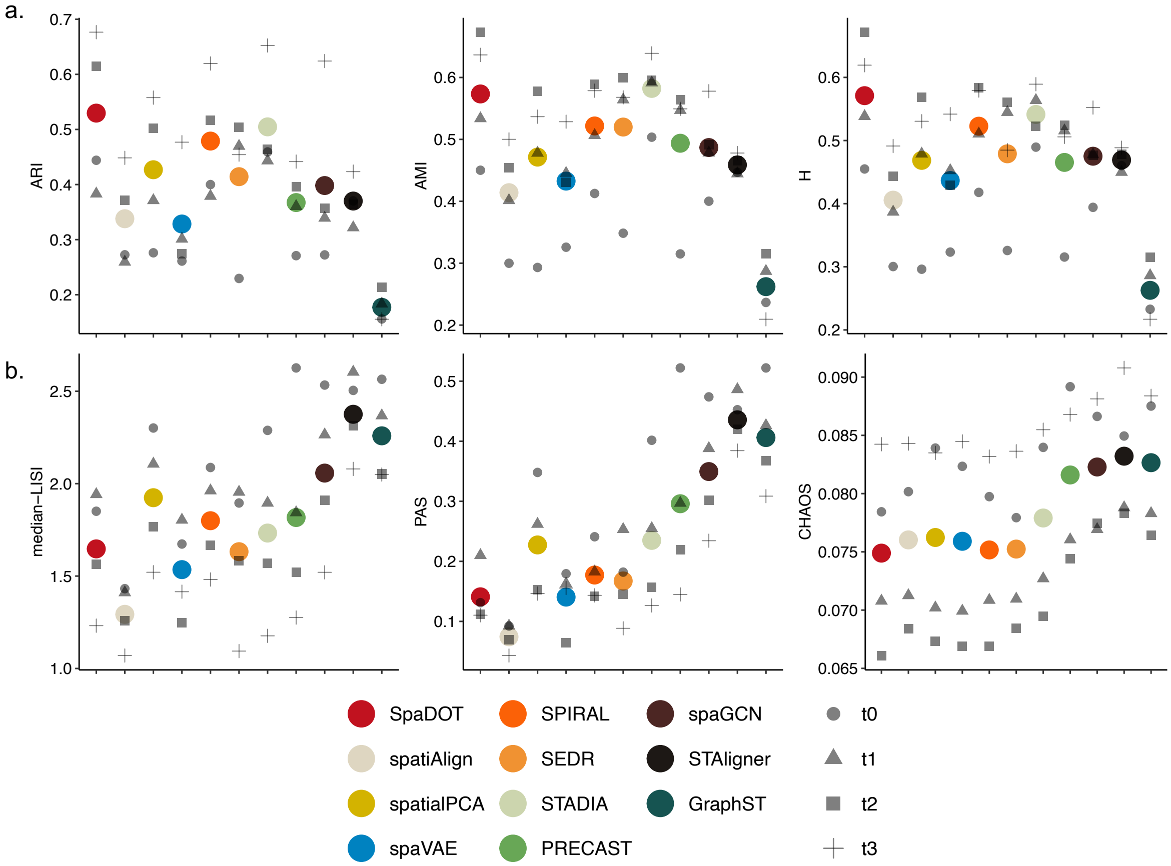


**Supplementary Figure S14.** a) Spatial domain detection accuracy evaluation using three metrics: adjusted Rand index (ARI), adjusted Mutual Information (AMI) and Homogeneity (H). Colored dots indicate the average performance across all time points, while grey dots with different shapes represent performance at individual time points. b) Spatial continuity evaluation using three metrics: median local inverse Simpson index (LISI), percentage of abnormal spots (PAS), and spatial CHAOS score (CHAOS). Colored dots indicate the average performance across all time points, while grey dots with different shapes represent performance at individual time points.

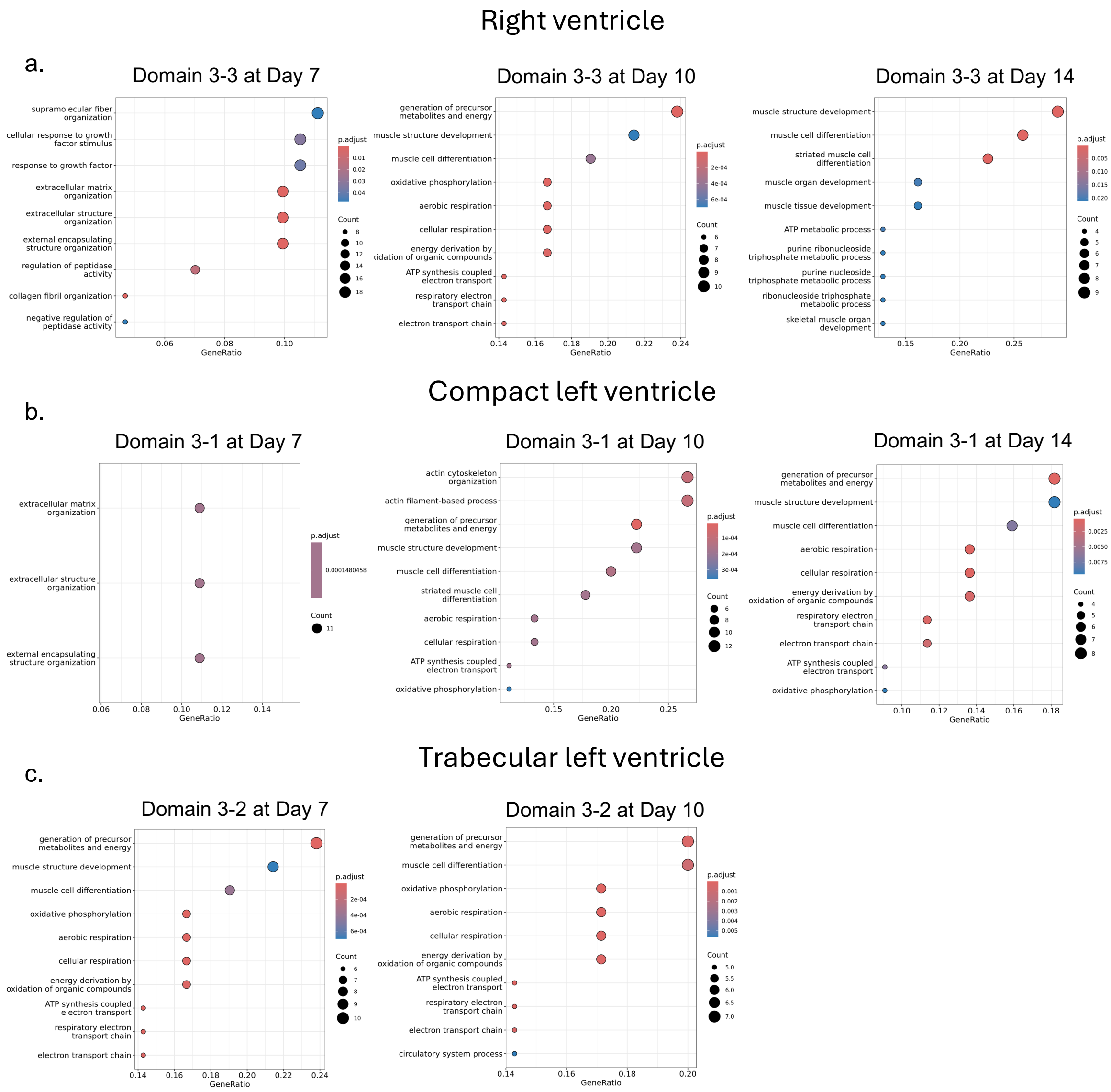


**Supplementary Figure S15.** a) Gene ontology (GO) enrichment for domains detected as right ventricle across all time points. b) Gene ontology (GO) enrichment for domains detected as compact left ventricle across all time points. c) Gene ontology (GO) enrichment for domains detected as trabecular left ventricle on Day 7 and Day 10.

a.

### Domain 5-1 vs Domain 5-2 UP

| Index | Name | P-value | Adjusted p-value | Odds Ratio | Combined score |
|---|---|---|---|---|---|
| 1 | Cardiomyocyte Heart Mouse | 7.738e-20 | 1.462e-17 | 303.01 | 13334.28 |
| 2 | Cardiomyocyte Heart Human | 1.010e-11 | 9.546e-10 | 96.46 | 2442.17 |
| 3 | Smooth Muscle Cell Lung Human | 0.000001366 | 0.00006454 | 248.93 | 3361.40 |
| 4 | Cardiomyocyte Heart Muscle Human | 0.000001366 | 0.00006454 | 248.93 | 3361.40 |
| 5 | Migration Phase Fetal Germ Cell Fetal Gonad Human | 0.000005616 | 0.0002123 | 11.52 | 139.26 |
| 6 | Cardiomyocyte Embryo Mouse | 0.00003712 | 0.001169 | 57.42 | 585.72 |
| 7 | Axin2-Palpha+ Cell Lung Mouse | 0.00006365 | 0.001719 | 46.64 | 450.67 |
| 8 | Cardiomyocyte Embryo Human | 0.0001688 | 0.003987 | 163.90 | 1423.80 |
| 9 | Myocyte Aorta Mouse | 0.0002525 | 0.005302 | 122.92 | 1018.29 |
| 10 | P53-like Cell Bladder Human | 0.0004688 | 0.008860 | 81.94 | 628.09 |

### Domain 5-1 vs Domain 5-2 DOWN

| Index | Name | P-value | Adjusted p-value | Odds Ratio | Combined score |
|---|---|---|---|---|---|
| 1 | Fibrocartilage Chondrocyte Articular Cartilage Human | 2.070e-20 | 1.186e-17 | 29.56 | 1339.69 |
| 2 | DCLK1+ Progenitor Cell Large Intestine Human | 1.485e-18 | 4.254e-16 | 30.00 | 1231.43 |
| 3 | Fibroblast Skin Mouse | 1.242e-17 | 2.373e-15 | 30.31 | 1180.06 |
| 4 | Medullary Cell Kidney Mouse | 5.296e-17 | 7.586e-15 | 18.45 | 691.35 |
| 5 | Fibroblast Heart Mouse | 7.727e-15 | 8.855e-13 | 74.92 | 2434.60 |
| 6 | Fibroblast Stomach Human | 1.419e-13 | 1.060e-11 | 51.85 | 1533.92 |
| 7 | Schwalie Et al.Nature.G2 Adipose Tissue Mouse | 1.430e-13 | 1.060e-11 | 13.89 | 410.74 |
| 8 | Leydig Precursor Cell Fetal Gonad Human | 1.479e-13 | 1.060e-11 | 12.53 | 370.12 |
| 9 | Fibroblast Undefined Mouse | 2.757e-13 | 1.755e-11 | 118.32 | 3421.84 |
| 10 | Fibroblast Lung Human | 7.888e-13 | 4.520e-11 | 42.12 | 1173.68 |

b.

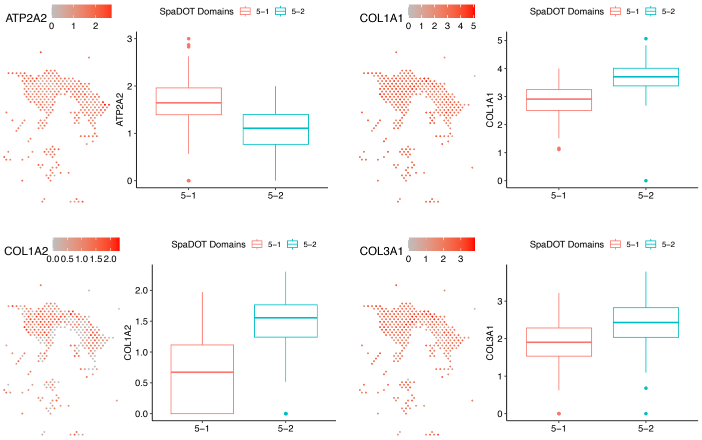


**Supplementary Figure S16.** a) Enrichment analysis using EnrichR (CellMarker 2024 category) based on valve related domain-specific up- and down-regulated genes at Day 14. b) Marker gene expression across valvular domains 5-1 and 5-2 at Day 14. *ATP2A2* is enriched in the atrioventricular valve, while *COL1A1*, *COL1A2*, and *COL3A1* are collagen fiber markers specific to semilunar valves. Boxplots display expression distributions of *ATP2A2*, *COL1A1*, *COL1A2*, and *COL3A1* across the predicted domains. The center line indicates the median; box limits represent the upper and lower quartiles; whiskers denote 1.5× the interquartile range.

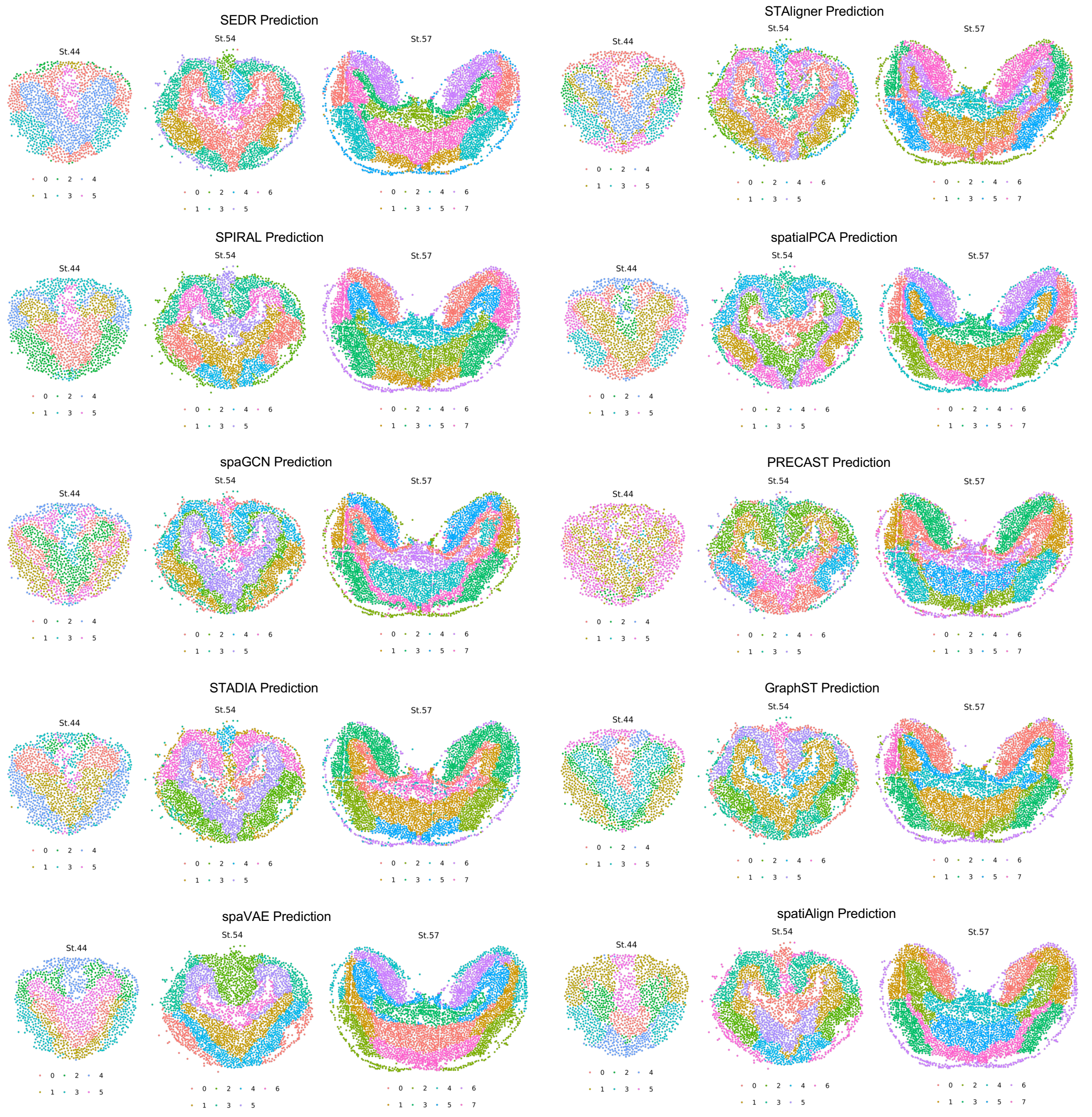


**Supplementary Figure S17.** Spatial domain layouts predicted by ten competing methods for comparison in developing axolotl telencephalon.

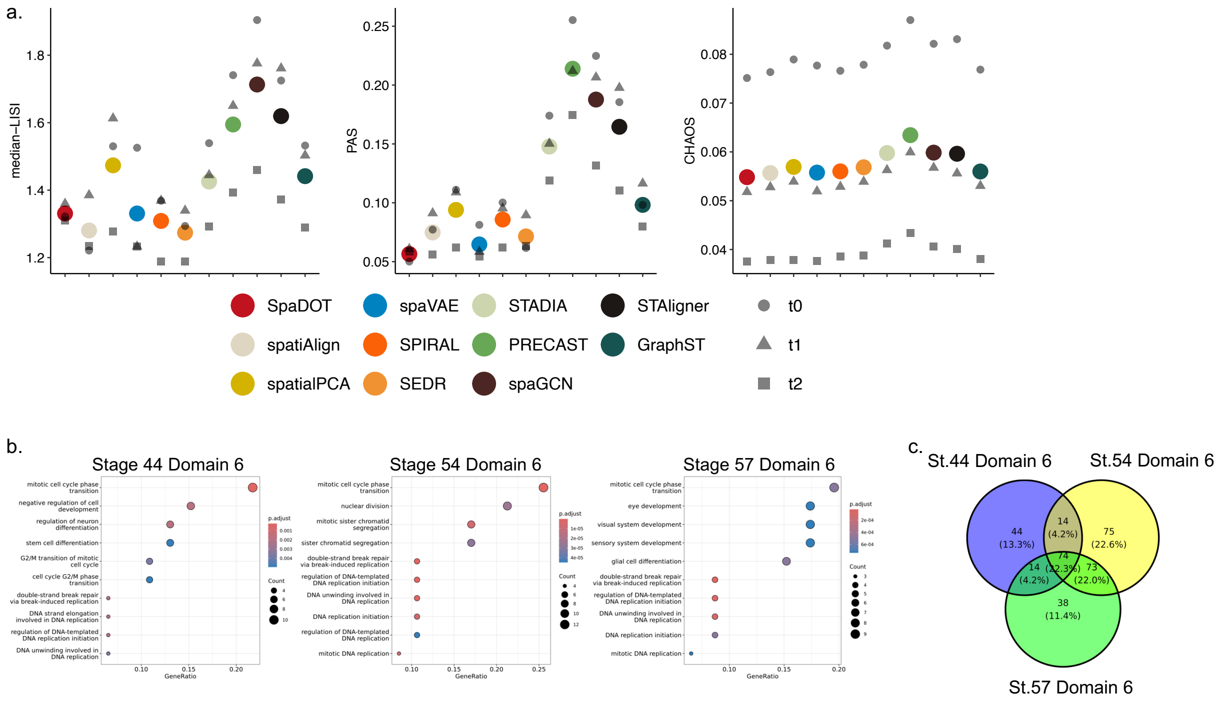


**Supplementary Figure S18.** a) Spatial continuity evaluation using three metrics: median local inverse Simpson index (LISI), percentage of abnormal spots (PAS), and spatial CHAOS score (CHAOS). Colored dots indicate the average performance across all time points, while grey dots with different shapes represent performance at individual time points. b) Gene ontology (GO) enrichment for domains detected as ventricular zone (domain 6) across all time points. c) Venn diagrams showing the number of overlapping GO terms among ventricular zone (domain 6) across all time points.

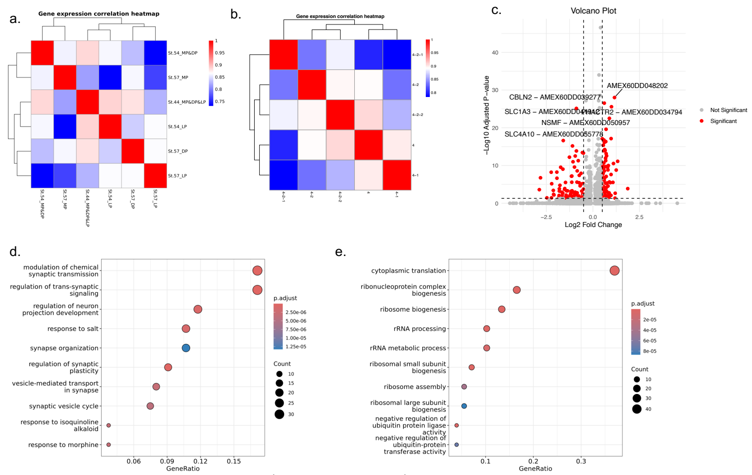


f.

Domain 4-2-1 vs Domain 4-2-2 UP

| Index | Name | P-value | Adjusted p-value | Odds Ratio | Combined score |
|---|---|---|---|---|---|
| 1 | Neuroendocrine Cell Trachea Mouse | 5.592e-10 | 1.549e-7 | 5.77 | 122.95 |
| 2 | Neuron Brain Mouse | 1.727e-9 | 2.393e-7 | 4.78 | 96.47 |
| 3 | Type I Spiral Ganglion Neuron Brain Mouse | 0.000004769 | 0.0004403 | 2.92 | 35.75 |
| 4 | Oligodendrocyte Brain Mouse | 0.00002810 | 0.001946 | 3.81 | 39.92 |
| 5 | Enteroendocrine Precursor Cell Intestinal Crypt Mouse | 0.00004187 | 0.002210 | 4.43 | 44.64 |
| 6 | Mucosal-associated Invariant T Cell Liver Human | 0.00005268 | 0.002210 | 10.11 | 99.59 |
| 7 | Neuron Epithelium Human | 0.00005586 | 0.002210 | 23.02 | 225.44 |
| 8 | Neuron Hypothalamus Mouse | 0.0004654 | 0.01523 | 23.79 | 182.52 |
| 9 | Delta Cell Pancreas Human | 0.0005611 | 0.01523 | 22.09 | 165.34 |
| 10 | Astrocyte Embryonic Prefrontal Cortex Human | 0.0005690 | 0.01523 | 3.51 | 26.20 |

Domain 4-2-1 vs Domain 4-2-2 DOWN

| Index | Name | P-value | Adjusted p-value | Odds Ratio | Combined score |
|---|---|---|---|---|---|
| 1 | Transit Amplifying (TA) Cell Intestinal Crypt Mouse | 2.634e-42 | 8.430e-40 | 65.91 | 6309.90 |
| 2 | Goblet Cell Intestinal Crypt Mouse | 6.410e-32 | 1.026e-29 | 26.05 | 1871.17 |
| 3 | Schwalie Et al.Nature.P1 Adipose Tissue Mouse | 1.384e-26 | 1.476e-24 | 30.92 | 1841.11 |
| 4 | Leydig Precursor Cell Fetal Gonad Human | 6.387e-24 | 5.109e-22 | 23.45 | 1252.43 |
| 5 | Microglial Cell Brain Human | 3.213e-22 | 2.057e-20 | 15.84 | 784.04 |
| 6 | Mitotic Fetal Germ Cell Fetal Gonad Human | 4.137e-22 | 2.207e-20 | 11.02 | 542.51 |
| 7 | Basal Cell Trachea Mouse | 8.621e-20 | 3.941e-18 | 36.11 | 1585.20 |
| 8 | Schwalie Et al.Nature.G4 Adipose Tissue Mouse | 3.298e-18 | 1.319e-16 | 21.81 | 877.91 |
| 9 | Astrocyte Embryonic Prefrontal Cortex Human | 1.029e-9 | 3.658e-8 | 8.24 | 170.58 |
| 10 | Schwalie Et al.Nature.G1 Adipose Tissue Mouse | 4.644e-9 | 1.486e-7 | 11.12 | 213.46 |

**Supplementary Figure S19.** a) Heatmap displaying Spearman correlation of averaged gene expression among ground truth domains annotated as medial pallium (MP), dorsal pallium (DP), and lateral pallium (LP) across all time points. b) Heatmap showing Spearman correlation of averaged gene expression among SpaDOT-predicted domains associated with MP, DP, and LP across all time points. c) Enrichment analysis using EnrichR (CellMarker 2024 category) based on domain-specific up- and down-regulated genes related to MP/DP at Stage 57.

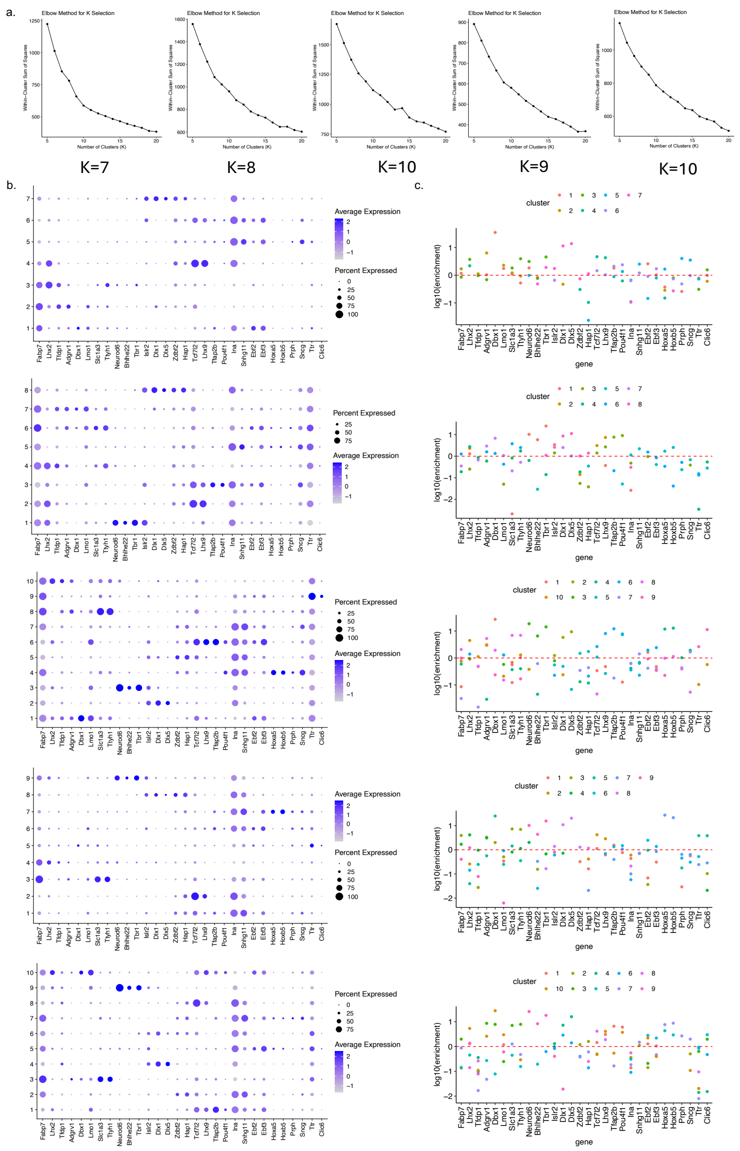


**Supplementary Figure S20.** a) Elbow plots showing the within-cluster sum of squares (WSS) for each time point. Elbow points are selected as 7, 8, 10, 9, and 10, based on a WSS difference threshold > 20 and then the highest WSS difference ratio. b) Dot plots showing marker gene enrichment for the uncurated, original domains predicted by SpaDOT. Dot color indicates enrichment level, and dot size represents the proportion of spots expressing each marker gene within the domain. c) Marker gene enrichment based on averaged expression in one domain versus all others predicted by SpaDOT. The y-axis shows log10-transformed enrichment values, with the red dashed line indicating a fold enrichment of 1.

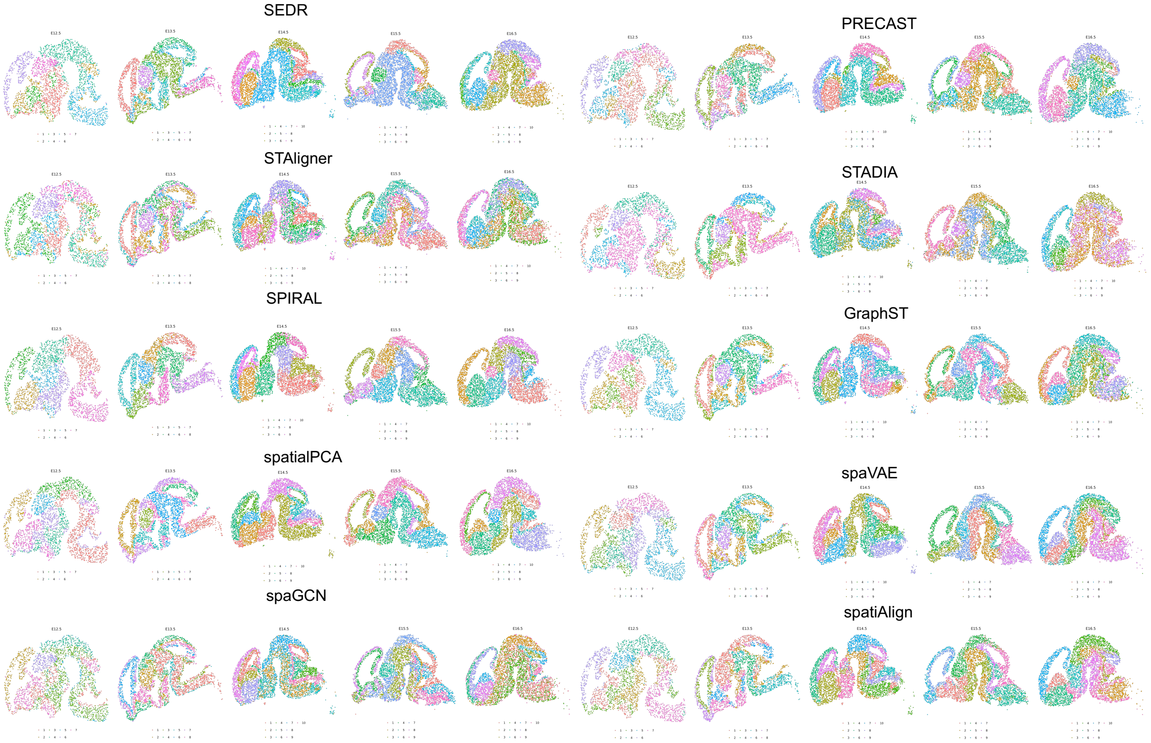


**Supplementary Figure S21.** Spatial domain layouts predicted by ten competing methods for comparison in developing mouse whole brain.

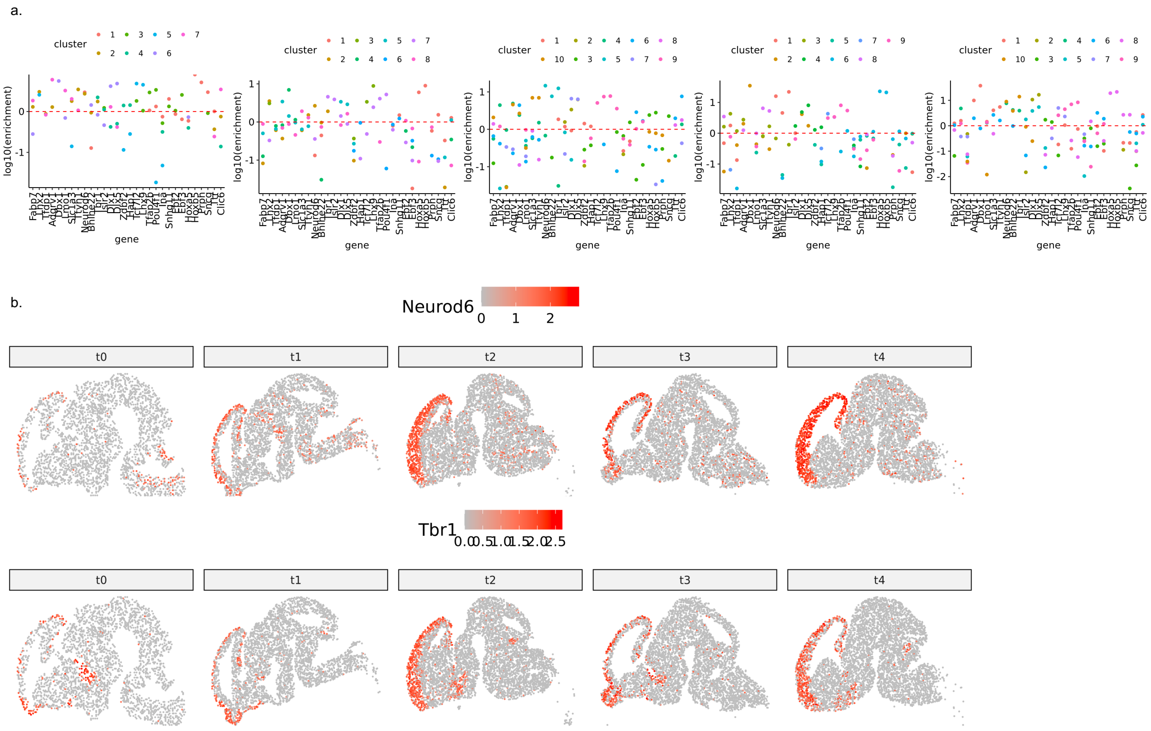


**Supplementary Figure S22.** a) Marker gene enrichment based on averaged expression in one domain versus all others predicted by SpatialPCA. The y-axis shows log10-transformed enrichment values, with the red dashed line indicating a fold enrichment of 1. b) Spatial expression patterns of key marker genes across all developmental stages. *Neurod6* and *Tbr1* are pallium markers.

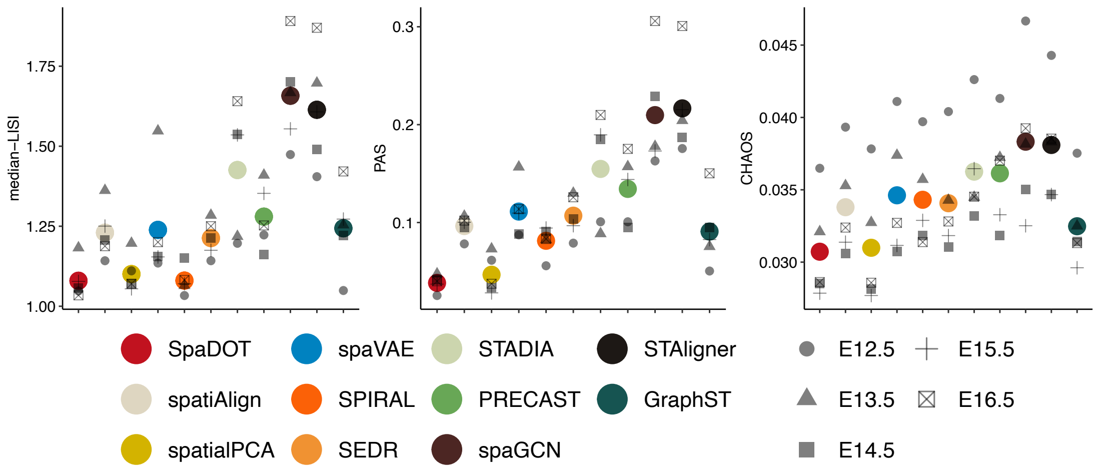


**Supplementary Figure S23.** Spatial continuity evaluation using three metrics: median local inverse Simpson index (LISI), percentage of abnormal spots (PAS), and spatial CHAOS score (CHAOS). Colored dots indicate the average performance across all time points, while grey dots with different shapes represent performance at individual time points.

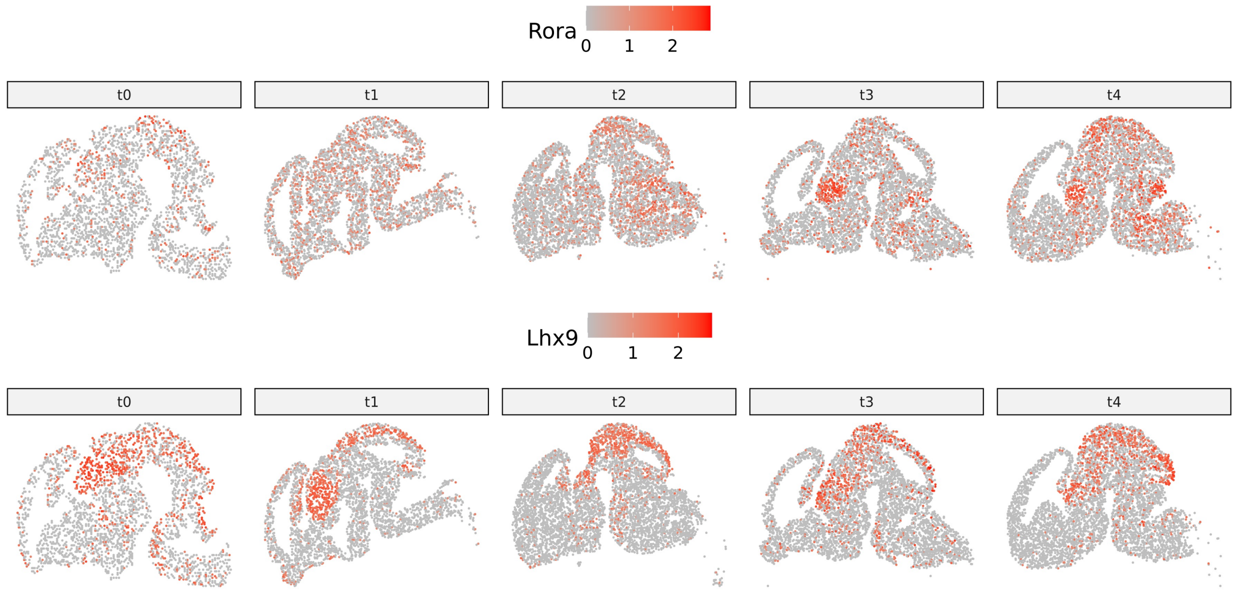


**Supplementary Figure S24.** Spatial expression patterns of key marker genes across all developmental stages. *Rora* and *Lhx9* are thalamus markers.
